# Calcium-triggered (de)ubiquitination events in synapses

**DOI:** 10.1101/2024.07.04.602026

**Authors:** Sofia Ainatzi, Svenja V. Kaufmann, Ivan Silbern, Svilen V. Georgiev, Sonja Lorenz, Silvio O. Rizzoli, Henning Urlaub

## Abstract

Neuronal communication relies on neurotransmitter release from synaptic vesicles (SVs), whose dynamics are controlled by calcium-dependent pathways, as many thoroughly studied phosphorylation cascades. However, little is known about other post-translational modifications, as ubiquitination. To address this, we analysed resting and stimulated synaptosomes (isolated synapses) by quantitative mass spectrometry. We identified more than 5,000 ubiquitination sites on ∼2,000 proteins, the majority of which participate in SV recycling processes. Several proteins showed significant changes in ubiquitination in response to calcium influx, with the most pronounced changes in CaMKIIα and the clathrin adaptor protein AP180. To validate this finding, we generated a CaMKIIα mutant lacking the ubiquitination target site (K291) and analysed it both in neurons and non-neuronal cells. K291 ubiquitination influences CaMKIIα activity and synaptic function by modulating its autophosphorylation at a functionally important site (T286). We suggest that ubiquitination in response to synaptic activity is an important regulator of synaptic function.

## Introduction

In a chemical synapse, information flows from a presynaptic neuron to a postsynaptic cell through the Ca^2+^-regulated release of neurotransmitters (NTs). NTs are stored in synaptic vesicles (SVs) located within the presynaptic nerve terminal. Functionally distinct pools of SVs co-exist within the presynaptic bouton, with only a small fraction actively participating in SV trafficking during low frequency stimulation (Denker, 2010). Specifically, SVs that are docked at specialized release sites, known as the active zone (AZ), are primed for SV exocytosis/fusion. The SV docking and priming at the AZ is mediated by a large protein complex consisting of scaffolding proteins (RIM, Munc13, RIM-BP, liprins, ELKS, bassoon and piccolo) (Südhof, 2012). Primed SVs at the AZ are the first to fuse with the presynaptic plasma membrane in response to stimulation, forming the readily releasable pool (RRP) (Denker, 2010). Exocytotic fusion of SVs is mediated by the SNAREs (Jahn and Fasshauer, 2012) and regulated by the Ca^2+^-sensing protein synaptotagmin (Sudhof, 2012) and other proteins, such as Munc18(Toonen and Verhage, 2007) and complexins (Sudhof, 2012). After SV exocytosis, SV membranes are endocytosed primarily by clathrin-mediated endocytosis (CME) (Chanaday et al., 2019) and SVs are regenerated to participate in another round of NT release.

Post-translational modifications (PTMs), such as protein phosphorylation, serve as a molecular mechanism for fine-tuning the SV cycle. Arguably, the best characterized example of such modulation is the regulation of SV mobility and availability for exocytosis by a conserved family of phosphoproteins, the synapsins (Ceccaldi et al., 1995; Greengard et al., 1994). Synapsins are SV-associated proteins that link SVs together to form a cluster of SVs, the reserve pool (RP) (Cesca et al., 2010; Denker, 2010). Several presynaptic kinases, including Ca^2+^/calmodulin-dependent kinase II (CaMKII), regulate the phosphorylation state of synapsins (Cesca et al., 2010). CaMKII-dependent phosphorylation of synapsin at specific sites (S566, S603) during stimulation reduces binding to actin and SVs, thereby releasing the SVs from the RP and making them available for exocytosis (Cesca et al., 2010). In contrast to synapsins, several proteins involved in CME, termed dephosphins (Cousin and Robinson, 2001), are dephosphorylated during stimulation to facilitate SV endocytosis, such as the adaptor proteins epsin, eps15 and AP180(Cousin and Robinson, 2001; Kohansal-Nodehi et al., 2016). In addition to these proteins, recent mass spectrometry (MS) based phosphoproteomic studies revealed that a large number of presynaptic proteins undergo phosphorylation changes in response to Ca^2+^ influx, suggesting that phosphorylation is a driver of the changes in protein–protein interactions that occur in the synapse during stimulation (Engholm-Keller et al., 2019; Kohansal-Nodehi et al., 2016; Silbern et al., 2021).

In addition to phosphorylation, a growing body of studies indicates that another PTM, protein ubiquitination, plays an important role at the synapse. In general, Κ48-linked ubiquitin chains target substrate proteins for degradation by the 26S proteasome (Kwon and Ciechanover, 2017), thereby regulating protein quality control as well as diverse other cellular functions ^30^, whereas monoubiquitination and other ubiquitin linkage types are often associated with non-degradative processes, such as membrane protein trafficking, DNA repair, signalling pathways, and the activation of protein kinases (Kwon and Ciechanover, 2017). Multiple monoubiquitination and K- 63 linkages have been implicated in the multivesicular endosomal sorting of plasma membrane proteins followed by their degradation in lysosomes (Raiborg and Stenmark, 2009). Ubiquitination typically occurs at the primary amino group of Lys residues of substrates, mediated by the sequential action of three enzymes (ubiquitin-activating enzyme (E1), ubiquitin-conjugating enzyme (E2), and ubiquitin ligase (E3) enzymes), and its removal is achieved by deubiquitinating enzymes (DUBs) (Kwon and Ciechanover, 2017).

Indeed, synaptic proteins undergo ubiquitination and degradation by the proteasome (UPS) (Chin et al., 2002; Lazarevic et al., 2011; Speese et al., 2003; van Roessel et al., 2004; Wheeler et al., 2002), including a number of AZ proteins, such as RIM and Munc13(Jiang et al., 2010; Lazarevic et al., 2011; Speese et al., 2003; Yao et al., 2007). Spatiotemporal regulation of ubiquitination and degradation is achieved at the presynaptic nerve terminals through the AZ scaffold proteins bassoon and piccolo (Waites et al., 2013). Lastly, perturbation of the UPS via pharmacological agents was shown to influence NT release in different preparations (Rinetti and Schweizer, 2010; Speese et al., 2003). Apart from the degradative roles of ubiquitin, ubiquitin also has non-degradative roles in the presynapse. One element suggesting a modulatory role of ubiquitination is the high rate at which protein ubiquitination occurs and its correlation with synaptic activity. Specifically, a rapid decrease in the total ubiquitination levels was observed upon Ca^2+^ influx in isolated nerve terminals, an effect that was only partially reversed by proteasome inhibition (Chen et al., 2003). The same study showed that the clathrin-adaptor proteins, epsin-1 and eps15, underwent fast deubiquitination in response to depolarization-triggered Ca^2+^ influx (Chen et al., 2003). Finally, acute pharmacological inhibition of protein ubiquitination in cultured neurons was shown to elicit a rapid increase in spontaneous neurotransmitter release (Rinetti and Schweizer, 2010).

However, it is not known which synaptic proteins are ubiquitinated at which sites and whether their ubiquitination status changes in different states of the synapse. Here, liquid chromatography-mass spectrometry (LC-MS/MS) offers the possibility of comprehensive identification and relative quantification of ubiquitination sites on proteins. Based on improved immunoaffinity purification strategies of ubiquitinated peptides (Kim et al., 2011; Xu et al., 2010), it has become evident that ubiquitination is a very prominent protein modification in cells, and importantly, that changes in the ubiquitination state of proteins are observed under different conditions in various cellular systems (Hansen et al., 2021; Kim et al., 2011; Povlsen et al., 2012; Sarraf et al., 2013; Satpathy et al., 2015; Udeshi et al., 2020; Wagner et al., 2011). To date, only a few studies have characterized protein ubiquitination in brain tissue by MS, but no ubiquitinome analyses have been performed in isolated nerve terminals (Na et al., 2012; Wagner et al., 2012). Particularly in this context, it remains to be determined which synaptic proteins undergo ubiquitination changes in response to Ca^2+^ influx, illuminating the non-degradative functions of ubiquitination in synapses. In this study, we performed a proteome-wide quantification analysis of ubiquitinated proteins in isolated nerve terminals, termed “synaptosomes”(Gray and Whittaker, 1962). Synaptosomes are considered a valid model system of the synapse, as they contain SVs, mitochondria and the molecular machinery required for the SV cycle (Gray and Whittaker, 1962; Nicholls et al., 1987). We isolated synaptosomes from rat brain and subjected them to a bottom-up proteomic workflow incorporating antibody-based enrichment of formerly ubiquitinated peptides, followed by labeling with isobaric tandem mass tag (TMT) reagents and LC-MS/MS analysis (Kim et al., 2011; Thompson et al., 2003). Our analysis identified 41 proteins that undergo significant ubiquitination changes in response to Ca^2+^ and provides the first synapse-specific inventory of ubiquitination sites. Prominent changes in ubiquitination are observed for Ca^2+^/calmodulin-dependent kinase II (CaMKII) and the clathrin adaptor protein 180 (AP180) upon depolarization. Lastly, we have functionally characterized one of these newly discovered ubiquitination sites, K291 of CaMKIIα.

## Results

### Description of the workflow and overview of results

In a first discovery approach, we performed a quantitative analysis of ubiquitinated proteins in chemically stimulated synaptosomes under Ca^2+^-restricted and Ca^2+^-rich conditions (**Figure 1**).

**Fig. 1:**
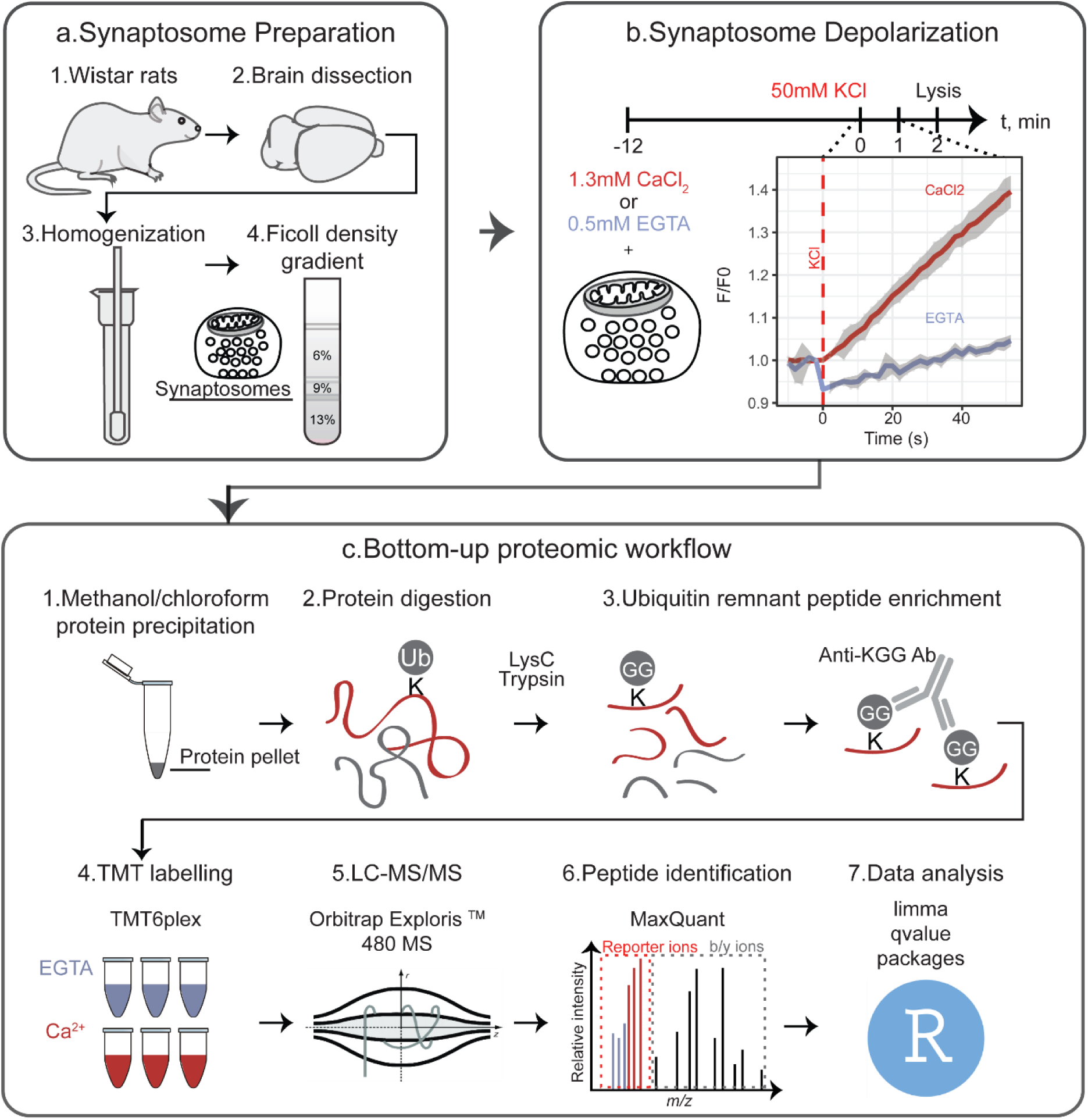
**Workflow for the quantitative analysis of ubiquitinated proteins in depolarized synaptosomes under different conditions**. (a) Synaptosomes were isolated from brains of 5–6-week-old Wistar rats by homogenisation of brain tissue followed by differential centrifugation and discontinuous Ficoll gradient centrifugation. (b) Synaptosome depolarization was induced by KCl in the presence of Ca^2+^ or the Ca^2+^- chelator EGTA and was quenched after two minutes by addition of lysis buffer. Three independent stimulations were performed for each condition (EGTA vs. Ca^2+^). (c) Equal amounts of proteins were subsequently precipitated by methanol/chloroform protein precipitation and sequentially digested with LysC and trypsin, followed by ubiquitin remnant-containing (K-ε-GG) peptide enrichment and chemical labelling with isobaric TMT6 reagents. Differently labelled peptides were combined and analysed by LC- MS/MS. Two independent TMT6 experiments were performed. Peptide identification and quantification was performed in MaxQuant and the extracted reporter ion intensities were further processed in R.

Membrane depolarization of synaptosomes was chemically induced by increasing the external concentration of KCl in the medium. To assess whether synaptosomes were responsive to chemical depolarization the release of glutamate was monitored upon stimulation as previously described (Nicholls et al., 1987) **(Figure S1a**). Chemical depolarization was applied for 2 min in a buffer containing either the Ca^2+^ chelator EGTA or Ca^2+^ (**Figure 1**). Subsequently, equal amounts of protein from differentially treated synaptosomes were precipitated and sequentially digested with LysC and trypsin. Tryptic digestion of ubiquitinated proteins leaves a ubiquitin-remnant di-glycine dipeptide on the lysine residue of the substrates (K-ε-GG) with a monoisotopic mass of 114.04 Da. We used commercially available antibodies that specifically recognize the ubiquitin remnant to enrich for K-ε-GG peptides. Subsequently, the K-ε-GG peptides were chemically labelled with TMT6 reagents, combined and analyzed by LC-MS/MS (**Figure 1**). In parallel, we compared the proteome of depolarized synaptosomes under Ca^2+^-restricted and Ca^2+^-rich conditions using TMT-based quantification approach, followed by off-line basic reversed phase (bRP) peptide fractionation and LC-MS/MS analysis.

Our analyses led to the identification and quantification of 5,258 confidently localized ubiquitination sites in more than 2,000 proteins, demonstrating that ubiquitination is a widespread post-translational modification of synaptic proteins covering a wide range of protein abundances in our sample (**Figure 2a**). Our data represent a unique data set of ubiquitination sites in the synapse. Many synaptic proteins were found highly ubiquitinated at more than 20 sites, such as synaptotagmin-1 (*Syt1*), Munc-18 (also known as *Stxbp1*) and CaMKIIα (**Figure 2a**). Comparison of our ubiquitination data set with a previous data set derived from whole mouse brain revealed more than 2,000 shared ubiquitination sites (Wagner et al., 2012) (**Figure 2d**). A similar comparative analysis of our data set with the PhosphositePlus ubiquitination data set (Hornbeck et al., 2015) showed that 65% of the ubiquitination sites identified by us have been previously reported in other studies. Our data set thus encompasses hitherto non-identified ubiquitination sites, such as four distinct sites in the C-terminal region of the AZ protein, RIM, a well-established substrate of ubiquitination (Yao et al., 2007) (**Figure S2a**), 12 novel sites mapped to the AZ protein piccolo (*Pclo*), four novel sites in complexin, and five novel sites mapped to the SNARE protein syntaxin-1 (*Stx1*) (**Suppl. Data 1**). Finally, the proteomic analysis resulted in the identification and quantification of approximately 5,800 unique proteins, hereafter referred to as the “synaptic proteome” of our sample, which was used as a true positive background in the pathway enrichment analysis (see below).

**Fig.2: Pathway.**
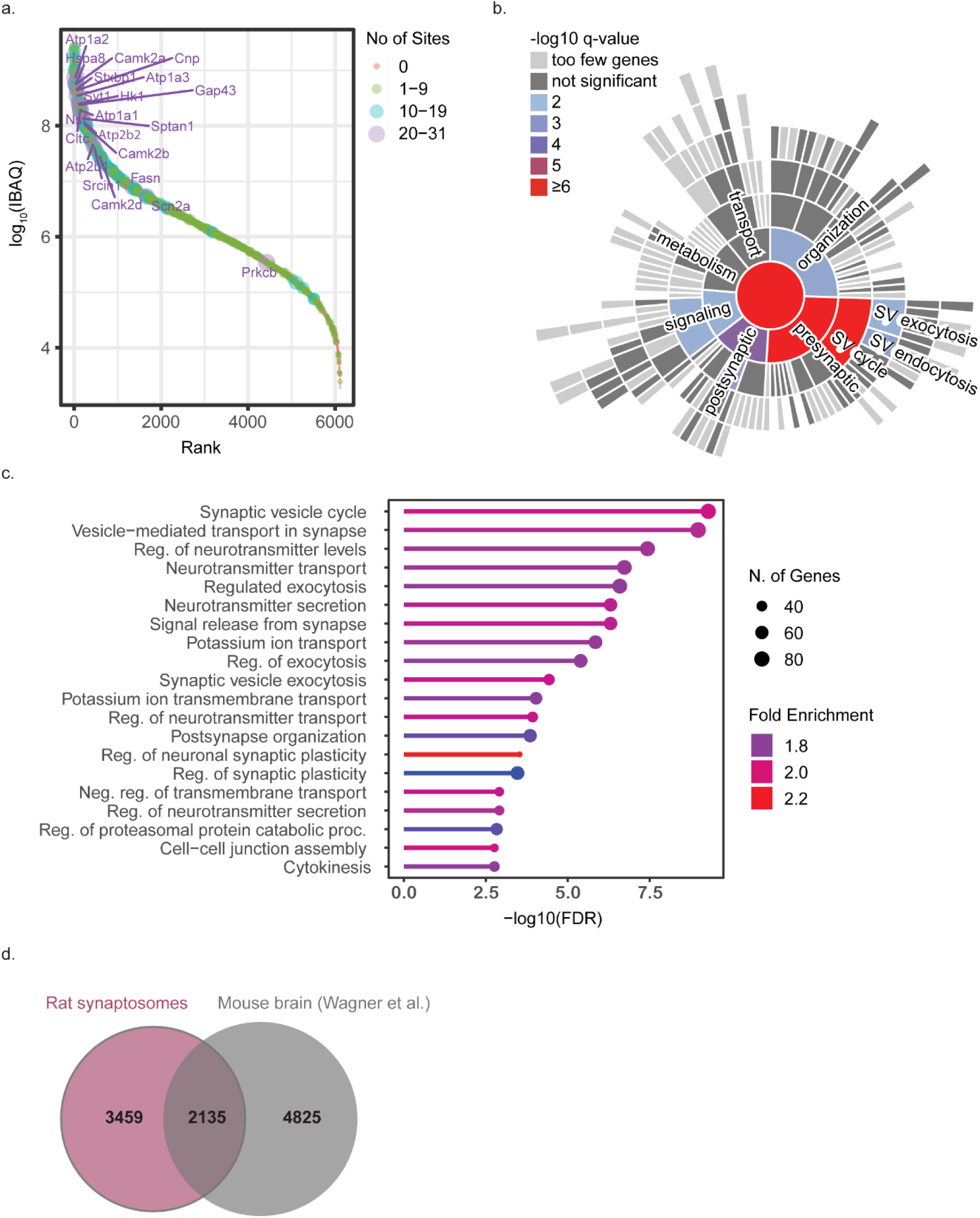
enrichment analysis of ubiquitinated proteins identified in synaptosomes and comparison of our data set with the literature. (a) Rank order of protein signals in the TMT experiment depicting the number of identified ubiquitination sites per protein. (b) Sunburst diagram depicting significantly enriched biological process terms based on the SynGo database (Koopmans et al., 2019). (c) Detailed list of enriched biological processes based on the ShinyGO (Ge et al., 2020). (d) Comparison of our ubiquitination data set derived from rat synaptosomes with a previous ubiquitination data set derived from mouse brain (Wagner et al., 2012) based on sequence similarity of the 6 amino acids flanking N- and C-terminal the modified lysine residue.

To obtain an overview of the functional properties of the identified ubiquitinated proteins we performed pathway enrichment analysis using SynGO, a synapse-specific database containing high quality annotations (Koopmans et al., 2019). Among the identified ubiquitinated proteins, 558 were mapped to unique annotated genes in the SynGO database (Koopmans et al., 2019). Subsequent enrichment analysis of these ubiquitinated proteins using our “synaptic proteome” as a custom background proteome revealed significantly enriched terms primarily associated with presynaptic functions such as the SV cycle, SV exo- and endocytosis (**Figure 2a,b**). Even though SynGO is a synapse-specific database with high quality annotations, it contains annotation exclusively for synaptic proteins. To obtain a more comprehensive view of the functional properties of ubiquitinated proteins, we performed a pathway enrichment analysis using ShinyGO, which uses annotations from the Ensembl database (Ge et al., 2020) (**Figure 2C**). Consistent with SynGO, enrichment analysis using ShinyGO revealed significantly enriched terms associated with synaptic functions including SV cycle, vesicle-mediated transport in the synapse, regulated exocytosis etc (**Figure 2c**). In addition to synaptic functions, ShinyGO enrichment analysis also revealed significantly enriched terms associated with the enzymatic machinery necessary for ubiquitination (**Figure 2c**). In particular, the term “regulation of proteasomal protein catabolic process” includes ubiquitin ligases as well as DUBs, which are of interest, as they may contribute to synapse-specific ubiquitination patterns (**Table S6, S7**). For a detailed list of enriched GO terms, see **Table S1, S2, S3**.

### Changes in protein ubiquitination in depolarized synaptosomes

Quantitative comparison of the ubiquitination sites of chemically depolarized synaptosomes under Ca^2+^-deprived and Ca^2+^-rich conditions demonstrated that only a small fraction of ubiquitination sites changed significantly in response to Ca^2+^ influx. Specifically, our quantitative analysis revealed 43 ubiquitination sites associated with 41 proteins that showed at least a 1.15-fold change at a false discovery rate (FDR) of 5% (**Figure 3a**). Both ubiquitination and deubiquitination events were observed, with deubiquitination events being slightly more pronounced. This is consistent with results of a previous study demonstrating a decrease in total ubiquitination levels in response to chemical depolarization (Chen et al., 2003). We note that we do not observe a change in the abundance of the ubiquitin chains of different linkage types upon stimulation, as reflected by the corresponding di-glycine modified remnants on ubiquitin itself.

**Fig. 3:**
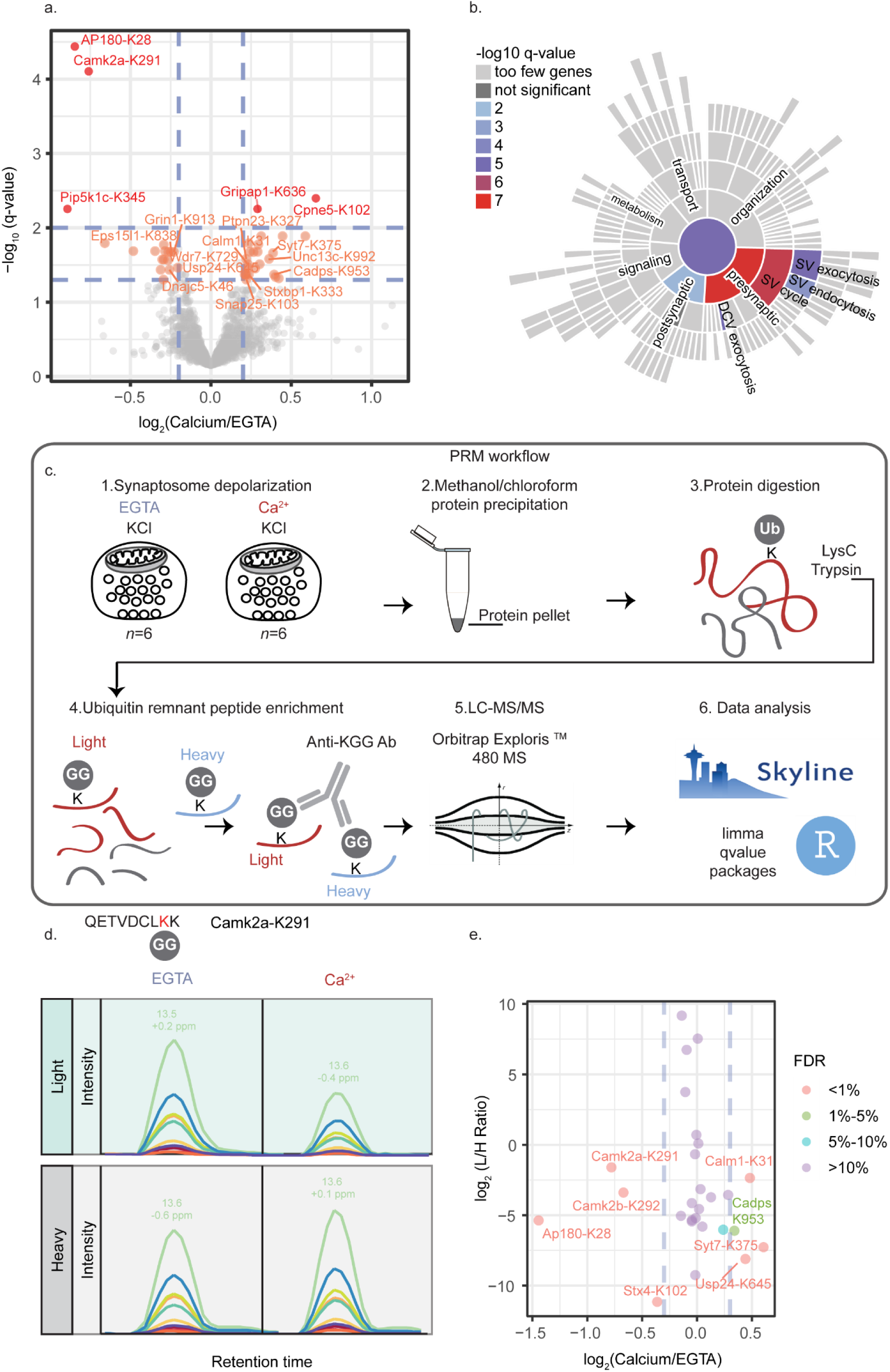
**Ubiquitination changes in depolarized synaptosomes under different stimuli**. (a) Volcano plot showing log2(intensity fold change) of ubiquitination sites quantified under Ca^2+^ vs. EGTA conditions against –log10(q-value). The colour encodes the significance of changes, highlighting with red and orange the ubiquitination sites that change significantly at FDRs of 1% and 5%, respectively. (b) Sunburst diagram depicting enriched biological process terms of proteins possessing regulated ubiquitination sites based on the SynGO database (Koopmans et al., 2019). (c) Synaptosome depolarization was induced by KCl in the presence of Ca^2+^ or the Ca^2+^-chelator EGTA and was quenched after two minutes by addition of lysis buffer. Six independent stimulations were performed for each condition (EGTA vs. Ca^2+^). Equal amounts of proteins were subsequently precipitated by methanol/chloroform protein precipitation and sequentially digested with LysC and trypsin. Standard/heavy peptides were spiked in the mixture of endogenous/light peptides prior to ubiquitin-remnant (K-ε-GG) peptide enrichment. Eluted (K-ε-GG) peptides were analysed by LC- MS/MS. Peptide identification and quantification was performed in Skyline and the extracted peak areas were further processed in R. (d) Examples of extracted fragment peak areas of endogenous/”light” and standard/”heavy” peptides corresponding to the ubiquitinated CaMKIIα peptide at K291, under calcium- deprived (EGTA) and calcium-free conditions. (e) Log2(light-to-heavy peptide intensity fold change) of ubiquitination sites quantified under Ca^2+^ vs. EGTA conditions against the log2 (light-to-heavy peptide intensity ratio) obtained by the PRM approach. The colour shows the statistical significance (FDR) of log2(light-to-heavy peptide intensity fold change).

To obtain an overview of the functional characteristics of proteins with regulated ubiquitination sites, we performed a pathway enrichment analysis using the SynGO database (Koopmans et al., 2019). Of the 41 proteins with regulated ubiquitination sites, only 18 were mapped to the SynGO database. Enrichment analysis with our “synaptic proteome” as a background revealed enriched terms associated primarily with presynaptic functions, namely, SV exo- and endocytosis as well as dense core vesicle exocytosis (**Figure 3b**). Specifically, proteins involved in SV endocytosis, such as AP180, EPS15L and Pip5k1c, were found to undergo deubiquitination in response to Ca^2+^ influx, with the clathrin adaptor protein AP180 showing the most prominent change (**Figure 3a**). Interestingly, the downregulated ubiquitination site of AP180 was mapped at K28, within the AP180 N-terminal homology (ANTH) domain (De Camilli et al., 2002; Stahelin et al., 2003) (**Figure S2b**). Like AP180, Ca^2+^/calmodulin-dependent kinase 2α (CaMKIIα) underwent a drastic decrease in its ubiquitination state in response to depolarization (**Figure 3a**). The downregulated ubiquitination site of CaMKIIα was mapped to K291 within the regulatory domain (i.e., the autoinhibitory domain) of CaMKIIα (**Figure S2c**). Conversely to endocytic proteins, active zone (AZ) proteins – such as Cadps1, Munc-18 (also known as *Stxbp1*) and synaptotagmin-7 (*Syt7*) – show enhanced ubiquitination upon stimulation (**Figure 3a, b**).

To validate differentially regulated ubiquitination sites, instead of “classical” orthogonal approaches (e.g. western blot analysis) we adopted a targeted MS approach, i.e. parallel reaction monitoring (PRM) (Peterson et al., 2012) (**Figure 3C**). PRM is the most sensitive and the least biased targeted assay for monitoring specific proteins/peptides of interest. It relies on synthetic isotope- labelled standard peptides that are added to the mixture of endogenous peptides in known amounts, thus allowing the (modified) endogenous peptide to be monitored quantitatively. Using synthetic isotope-labelled and modified peptides with the ubiquitin remnant at the lysine position, we targeted in total 9 regulated and 17 non-regulated ubiquitination sites according to the TMT experiment (**Table S4, Suppl. Data 2.1, 2.2**), including K-ε-GG peptides derived from ubiquitin itself (**Figure 3C**). Our analysis validates 6 out of 9 selected regulated sites, and 14 out of the 17 non- regulated ubiquitination sites, demonstrating the capacity of PRM as an additional validation strategy to reveal false positive and false negative hits (**Figure 3e, Suppl. Data 3**). Importantly, we confirmed the stark deubiquitination events on K28 of AP180 and K291 of CaMKIIα upon stimulation. Specifically, the ubiquitination of AP180 showed a decrease by a factor of 2.8, whereas the ubiquitination on CaMKIIα showed a decrease by a factor of 1.7. We do not attribute these fast and drastic changes in the ubiquitination pattern that we (and others (Chen et al., 2003)) observe to protein degradation for several reasons: first, we (and others) observe a rapid (within seconds to 2 min) deubiquitination upon depolarization of synapses; second, we do not observe a change in the abundance of the corresponding proteins, AP180 and CaMKIIα (**Figure S1b**); and third, AP180 and CaMKIIα are unusually long-lived, with average respective lifetimes of 52 and 16 days in cortex synaptosomes (Fornasiero et al., 2018).

### The regulatory region of CaMKIIα is a hotspot of PTMs

Our quantitative analysis revealed that CaMKIIα undergoes marked deubiquitination at K291 in response to stimulation, an observation that was further validated by our PRM analysis. This finding is interesting for several reasons: First, the ubiquitination site (K291) resides in the autoinhibitory domain of CaMKIIα, which is critical for regulating the enzyme’s activity (Colbran et al., 1988; Payne et al., 1988) (**Figure 4a**). Second, K291 is located close to a regulatory autophosphorylation site of CaMKIIα, T286, which when phosphorylated confers Ca^2+^- independent activity to the enzyme (Schworer et al., 1988; Thiel et al., 1988). Third, sequence comparison reveals that K291 is highly conserved among metazoan from cnidarians to human, as well as conserved within the CaMKIIα paralogue genes α, β and δ, suggesting its functional importance (**Figure S3a**). Lastly, the deubiquitination of CaMKIIα at K291 is reversible upon Ca^2+^- chelation, indicating a potential modulatory role of ubiquitin (**Figure S3c**).

**Fig 4:**
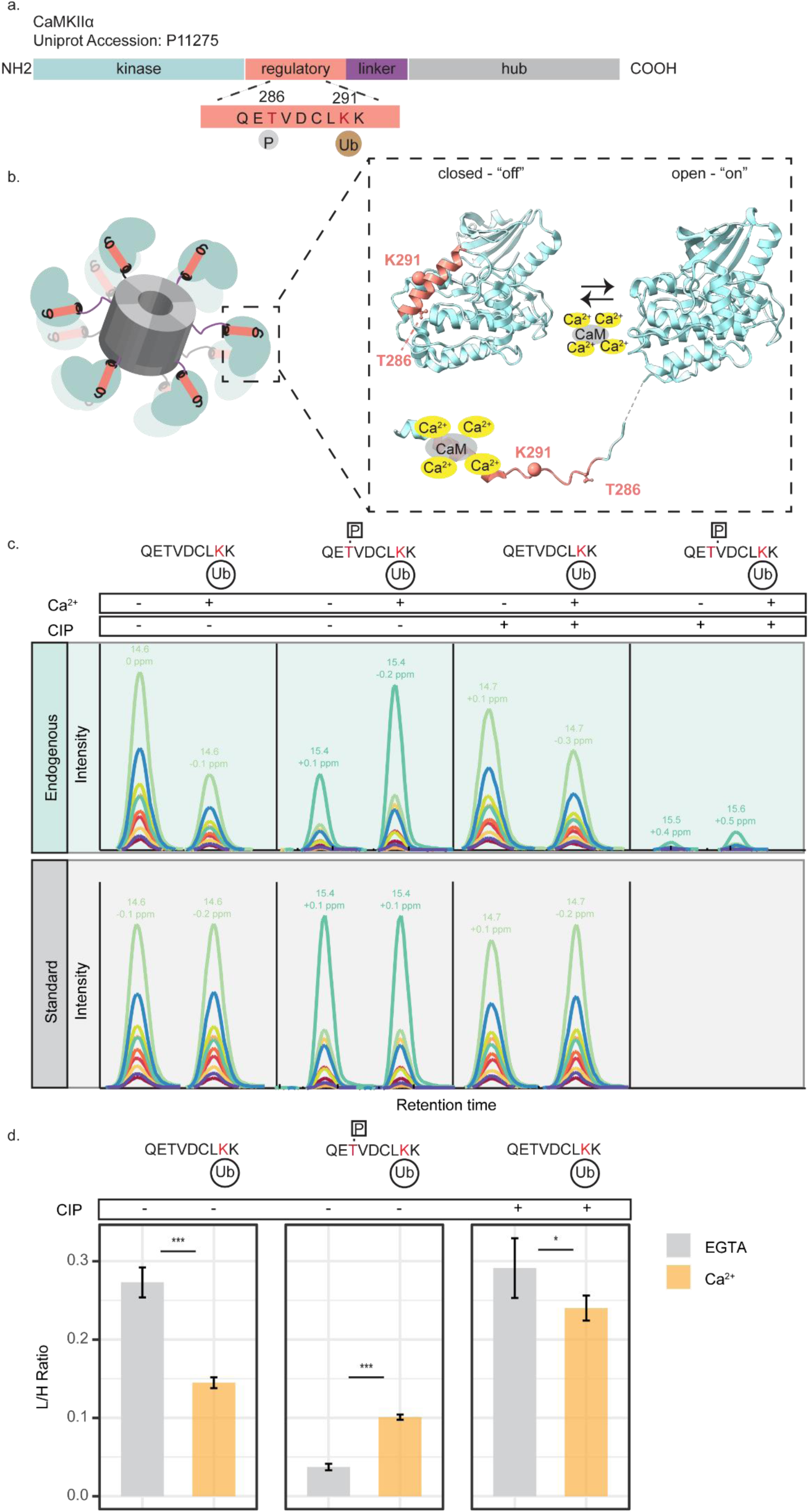
PTMs on the regulatory domain of Ca^2+^/calmodulin dependent kinase II α (CaMKIIα) and their quantification in depolarized synaptosomes under different conditions. (a) A horizontal bar represents the CaMKIIα sequence, with coloured regions showing the domains annotated according to Chao *et al*., 2011(Chao et al., 2011). The regulatory ubiquitination site (K291) resides in the regulatory domain of CaMKIIα very close to the autophosphorylation site (T286). (b) Dodecameric structure of CaMKIIα and the conformational states of a CaMKIIα subunit; in the close conformation (PDB code 2VN9) the regulatory domain folds back to the kinase domain, blocking access to the active site of the enzyme. Binding of Ca^2+^/calmodulin to the regulatory segment releases the active site of the enzyme, rendering the enzyme catalytically active and T286 accessible for phosphorylation (PDB code 2WEL). Part of K291 structure represented by a sphere is missing in the PDB codes. The PDB codes correspond to the human CaMKIIδ subunit 0-310 aa, which shares 92.58% sequence identity with human CaMKIIα and therefore we can safely assume that these domains have the same structure. (c) Examples of extracted traces of the product ions for endogenous/”light” and standard/”heavy” peptides corresponding to the ubiquitinated CaMKIIα peptide at K291 and the doubly modified CaMKIIα peptide at K291 and T286, before and after QuickCIP treatment. (d) Summary barplot showing the mean light-to-heavy peak area ratios. Limma statistical testing was performed to determine significant differences and account for the synaptosome preparation batch effect (N = six independent stimulation experiments, with two MS measurement replicates for each experiment). *P<0.05, ***P<0.001. We note that for the sake of simplicity we show here only one synaptosome preparation batch with three independent stimulation experiments. For a detailed view of both synaptosome batches refer to Figure S3.

It is well established that CaMKIIα undergoes activation upon Ca^2+^-influx,, exposing T286 for trans autophosphorylation by neighbouring CaMKIIα subunits of the multimeric structure (Hanson et al., 1994) (**Figure 4b**). Accordingly, CaMKIIα was reported to undergo phosphorylation at T286 in depolarized synaptosomes upon Ca^2+^ influx (Silbern et al., 2021). Importantly, the regulatory autophosphorylation and ubiquitination sites, T286 and K291 respectively, are located within the same tryptic peptide that was analysed by MS. Given the reported phosphorylation changes at T286, we hypothesized that the observed change in ubiquitination might be influenced by concomitant protein phosphorylation within the same peptide. To investigate this, we re-searched our quantitative data using both ubiquitin remnant (K-ε-GG) on lysine residues and phosphorylation on threonine residues as variable modifications. Indeed, the analysis led to the identification of the regulatory segment of CaMKIIα modified by both phosphorylation and ubiquitination at T286 and K291, respectively. While the ubiquitinated peptide species showed a ∼ 1.7-fold decrease upon depolarization of synapses, the doubly modified species showed a ∼2.2- fold increase in response to depolarization (**Figure S3d**). On this basis, we revised our ubiquitination analysis to consider that the measured decrease in CaMKIIα ubiquitination at K291 observed in the presence of Ca^2+^ may be caused (at least in part) by the Ca^2+^-induced autophosphorylation of CaMKIIα at T286. This is a prime example of the influence of two different posttranslational modifications within a given peptide in quantitative MS analyses, and it should also be considered in other cases (i.e., when comparing synaptic proteins in resting and depolarised synapses; see Discussion).

To determine more accurately the change of direction in CaMKIIα ubiquitination, we employed an absolute quantification approach. Specifically, we quantified the absolute amounts of the endogenous peptides of interest (**Table S5 & Suppl. Data 2.3, 2.4**), namely the singly ubiquitinated peptide and the doubly modified peptide of CaMKIIα, by PRM with known and equimolar amounts of stable isotope-labelled peptide standards in synaptosomal digests, followed by K-ε-GG peptide enrichment and PRM analysis. The analysis showed that upon stimulation the absolute levels of the doubly modified species increased from a light-to-heavy ratio of 0.034 to 0.1 (**Figure 4c,d, Figure S3e, Suppl. Data 4**), while the PRM signal of singly ubiquitinated peptide decreased from a light-to-heavy ratio of 0.27 to 0.14 (**Figure 4c,d, Figure S3e, Suppl. Data 4**). These PRM data demonstrate that deubiquitination of the singly ubiquitinated peptide is stronger than the increase in the doubly modified peptide through phosphorylation. We thus conclude that the observed apparent decrease in CaMKIIα ubiquitination upon stimulation of synaptosomes is, indeed, due to deubiquitination on K291.

To verify this finding, we further eliminated the confounding factor of T286 phosphorylation by treating the K-GG enriched peptides with phosphatase and repeating the absolute quantification of the singly ubiquitinated peptide of CaMKIIα. Indeed, after a QuickCIP treatment the doubly modified peptide was almost depleted (**Figure 4c**). In comparison, the levels of the singly ubiquitinated peptide decreased by a factor of 1.28 upon stimulation (**Figure 4c,d, Figure 3Se, Suppl. Data 4**).

### Ubiquitination of the CaMKIIα regulatory domain attenuates CaMKIIα T286 autophosphorylation and synaptic function

Our quantitative MS analysis demonstrate that the regulatory domain of CaMKIIα is a target for ubiquitination upon Ca^2+^ influx in depolarized synaptosomes. Given the fast response, the location and conservation of this site, we hypothesized that (de)ubiquitination might be involved in regulating CaMKIIα activity.

First, we investigated whether CaMKIIα stably expressed in HeLa cells can be used to assess the effect of K291 ubiquitination on CaMKIIα activity (**Figure 5b**), while also monitoring the quantitative changes of ubiquitination at K291 and autophosphorylation at T286 in response to Ca^2+^- influx using PRM-MS (**Table S5**). Ca^2+^ influx was induced by treating HeLa cells with the Ca^2+^ ionophore, ionomycin, and 1.8 mM of Ca^2+^ for 7 minutes. As expected, we observed an increase in T286 autophosphorylation upon stimulation (**Figure 5d, Suppl. Data 5**), indicative of CaMKIIα activation. In addition, we observed an increase in K291 ubiquitination on singly modified peptide and on doubly modified ones, bearing both phosphorylation at T286 and ubiquitination at K291 (**Figure 5c, Suppl. Data 5**). Together, these PRM data show that CaMKIIα undergoes phosphorylation and ubiquitination on T286 and K291, respectively, also in non-neuronal cells. However, we observe opposing changes in its ubiquitination state in the two systems we studied; whereas in CaMKIIα undergoes deubiquitination at K291 in response to Ca^2+^ influx in synaptosomes, the fluorescence-labelled CaMKIIα undergoes ubiquitination at K291 upon Ca^2+^ stimulation in HeLa cells.

**Fig. 5:**
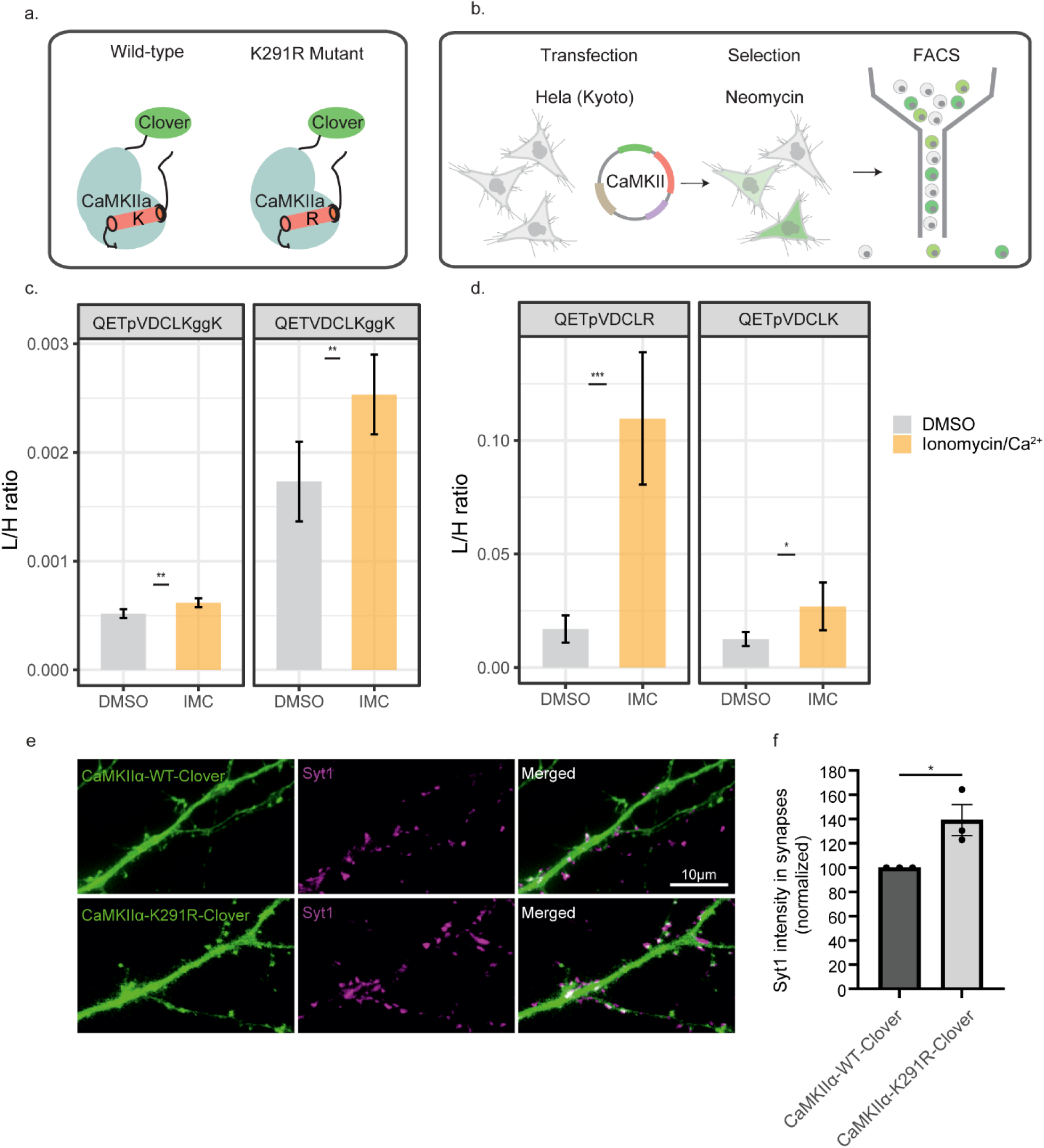
Functional assay to monitor the effects of CaMKIIα expression in HeLa cells and neurons. a) Generation of CaMKIIα K291R mutant that cannot be ubiquitinated at K291. b) Generation of HeLa Kyoto cell lines stably expressing either CaMKIIα-WT or the mutant variant K291R. c,d) Barplots illustrating the mean endogenous/”light”-to-standard/”heavy” peptide intensity in HeLa cells under different conditions (Ionomycin/Ca^2+^-vs-DMSO). Error bars correspond to the standard deviation. We note that we used the same standard/”heavy” peptide with the sequence QETpVDCLK to normalize the endogenous/light peptides QETpVDCLK and QETpVDCLR. A two-sample t-test was performed to determine significant differences (N = three independent stimulation experiments, with two MS measurement replicates for each experiment). *P<0.05, ***P<0.001. e) Neurons were transfected with either the wild type (WT) or the K291R variants of CaMKIIα and were analyzed by fluorescence microscopy 6-8 days later. The green channel indicates the CaMKIIα expression (Camui construct fluorescence), while the magenta channel shows anti-synaptotagmin 1 antibodies (directly conjugated to the fluorophore Atto647N), which are taken up by recycling synaptic vesicles, during a 60-minute incubation. After washing with Tyrode’s solution, the cells were fixed with PFA and imaged. f) Synapses were identified based on the synaptotagmin 1 signal, which was correlated with the Camui expression signal within the area of each synapse, using a Pearson correlation analysis. Subsequently, the fluorescence intensity of the synaptotagmin 1 label was quantified in the boutons in which the two signals were well correlated (meaning true presynaptic boutons, and not presynapses of non-transfected neurons that overlapped with CaMKIIα-expressing dendrites). A paired t-test between the wild type and the mutant was performed to determine significant differences (p=0.03, N = three independent experiments, with hundreds of synapses analysed for each experiment).

Despite this difference, we investigated the effect of the ubiquitination-deficient variant in the HeLa cells in response to ionomycin treatment, monitoring Ca^2+^-induced differences in T286 autophosphorylation compared to the wild type by PRM-MS. Strikingly, we observed that the K291R variant displayed significantly increased levels of T286 phosphorylation compared to the wild type (**Figure 5d, Suppl. Data 5**). This reveals that the absence of ubiquitination at K291 strongly increases the level of T286 for autophosphorylation, indicative of higher CaMKIIα activity.

In isolated synapses, CaMKII activity has been shown to correlate with synaptic activity, with high CaMKII activity being associated with increased rates of neurotransmitter release (Nichols et al., 1990). Therefore, we hypothesized that an increase in activity of the mutated, non-ubiquitinated CaMKII may lead to an increased an increased exocytosis of synaptic vesicles (SVs) and neurotransmitter release. To investigate this idea, we expressed the wild type and the K291R variant in cultured hippocampal neurons. We probed the neurons with an antibody directed against the lumenal (intravesicular) domain of synaptotagmin 1, which is taken up by recycling SVs and therefore provides an overall view of the changes in exocytosis and synaptic activity levels (Matteoli et al., 1992). We observed that synaptic boutons in the neurons expressing the K291R variant showed significantly higher synaptotagmin 1 labelling, which reflects an enhanced activity in exocytosis (**Figure 5 e,f**). As CaMKIIα activity correlates with exocytosis/neurotransmitter release, these results corroborate our hypothesis that the (de)ubiquitination at K291 influences CaMKIIα activity.

## Discussion

In this study, we used a quantitative proteomics strategy to characterise ubiquitination changes of synaptosomal proteins that occur upon Ca^2+^ influx in isolated nerve terminals. We generated an inventory of ubiquitination sites mapped to proteins in the synapse. 65% of the sites we identified were previously reported in PhosphositePlus database (Hornbeck et al., 2015), but not specifically for the synapse; 35% have not been reported before. Our inventory thus provides a rich resource for the neuroscience community, with many new ubiquitination sites that can be further analyzed functionally, e.g., interrogating their effects on protein–protein interaction relevant to SV exocytosis, endocytosis, and recycling during depolarization in the synapse. For instance, we have identified multiple novel ubiquitination sites in the active zone protein RIM1, a known substrate of ubiquitination (Yao et al., 2007). Interestingly, all four ubiquitination sites were mapped to the C-terminus of RIM1, a region known to interact with the substrate recognition subunit SCRAPPER (or *Fbxl20*) of the SKP1-CUL1-F-box E3 ligase (Yao et al., 2007). In addition, our dataset includes ubiquitination sites on 11 E3 ligases and 29 DUBs, potentially representing the E3 ligases and DUBs that may be particular relevant in the synapse (detailed list in Table S6 and S7).

Several conclusions can be drawn from our quantitative analysis of ubiquitination sites in depolarized synaptosomes under Ca^2+^-rich and Ca^2+^-depleted conditions. First, compared to protein phosphorylation, for which substantial changes were observed in the phosphoproteome (Silbern et al., 2021), a rather small fraction of proteins and their respective ubiquitination sites changed significantly 2 min after depolarization. Among others, regulated ubiquitination sites the levels of which increased during Ca^2+^ influx were mapped to the active- zone proteins, Cadps1, Munc-18 and Syt-7. In contrast, de-ubiquitination upon depolarization was observed for particular sites in proteins involved in clathrin-mediated endocytosis (CME), namely AP180, EPS15L and Pip5k1c. Stimulation-dependent deubiquitination of certain CME proteins, epsin-1 and EPS15, has been observed before in depolarized synaptosomes using immunoblotting (Chen et al., 2003). Although our study shows a similar effect on the ubiquitination state of CME proteins, we did not monitor deubiquitination of epsin-1 and ESP15, which may be attributed to the different stimulation times used in our study.

The deubiquitination of CME proteins is of particularly interest, as it coincides with their dephosphorylation that was previously observed in response to stimulation (Cousin and Robinson, 2001; Kohansal-Nodehi et al., 2016). Among the CME proteins observed to be dephosphorylated and deubiquitinated in response to stimulation, we detected the clathrin adaptor protein AP180. Dephosphorylation of AP180, which is known to take place in depolarized synaptosomes (Silbern et al., 2021), promotes its interaction with the AP-2 adaptor complex that is necessary for SV endocytosis (Hao et al., 1999). In this study, we report that, upon stimulation, AP180 undergoes stark deubiquitination at K28, which is located in the N-terminal ANTH domain known to bind to plasma-membrane regions containing phosphatidiylinositol-4,5-biphosphate (Ptdins(4,5)P2) (Ford et al., 2001; Stahelin et al., 2003). Specifically, K28 of the ANTH domain was shown to interact with the 5-phosphate of PtdIns(4,5)P2, together with two other lysine residues (K38, K40) and a histidine residue (H41) (Ford et al., 2001). We hypothesize that under resting conditions the ANTH domain is ubiquitinated at K28, which attenuates its interaction with PtdIns(4,5)P2-containing regions. Upon stimulation, AP180 undergoes deubiquitination at K28, resulting in its recruitment of AP180 to those regions where it can perform clathrin-adaptor functions. We propose that stimulation- dependent deubiquitination of AP180 allows its recruitment to endocytic regions, consistent with an increased rate of endocytosis in depolarized synaptosomes. Further investigation is required to establish the role of K28 ubiquitination in regulating the localization of the adaptor-clathrin protein AP180 and thus the rate of clathrin-mediated endocytosis. As the regulated ubiquitination and phosphorylation sites of AP180 do not reside within the same tryptic peptide, it is also unclear whether the modifications affect the same protein molecule.

In CaMKIIα, the regulated ubiquitination site (K291) was mapped to the autoinhibitory domain spanning the amino acid 274-314. This domain is subject to a number of other post-translational modifications (PTMs), including T286 autophosphorylation, S280 O-linked glycosylation and M281/282 oxidation (Erickson et al., 2013, 2008; Schworer et al., 1988; Thiel et al., 1988). In particular autophosphorylation at T286 occurs after Ca^2+^/calmodulin-dependent activation of CaMKIIα, rendering CaMKIIα Ca^2+^-independent (Schworer et al., 1988; Thiel et al., 1988). Similar to T286 autophosphorylation, other reported PTMs in the regulatory domain (Erickson et al., 2013, 2011, 2008; Hanson et al., 1994) occur after its Ca^2+^-triggered activation and render CaMKII Ca^2+^- independent. Our quantitative ubiquitinomic study revealed strong deubiquitination at K291 of CaMKIIα. Importantly, this deubiquination is accompanied by a stark phosphorylation at T286 upon Ca^2+^ influx, with both modifications co-existing within the same peptide and hence in the same protein. This provides an intriguing example of the tight crosstalk of phosphorylation and ubiquitination, a regulatory paradigm in eukaryotic cell biology. By employing absolute quantification and eliminating T286 autophosphorylation, we showed that the apparent CaMKIIα deubiquitination at K291 upon Ca^2+^ influx cannot be explained solely by the Ca^2+^-induced T286 autophosphorylation. We rather conclude that CaMKIIα undergoes deubiquitination at K291 in depolarized synaptosomes upon Ca^2+^ influx.

In contrast to depolarized synaptosomes, our PRM-MS in Hela cell culture expressing a CaMKIIα chimeric protein monitored an increase in K291 ubiquitination of CaMKIIα in response to Ca^2+^ influx. We attribute the opposing changes in ubiquitination to different E3 ligases and/or DUBs operating in these different biological systems. Importantly, a Lys-to-Arg substitution at residue 291 resulted in elevated levels of T286 autophosphorylation upon Ca^2+^ influx, suggesting that ubiquitination at K291 may affect CaMKIIα activity. In isolated synaptosomes we observed deubiquitination along with increased autophosphorylation in the regulatory domain of CaMKIIα upon Ca^2+^ influx, which led us to hypothesize that K291 ubiquitination may hinder autophosphorylation at T286. In line with this hypothesis, the expression of the non-ubiquitinated K291R variant in primary neurons resulted in enhanced synaptic activity compared to the wild-type form. Together these findings highlight the functional importance of K291 ubiquitination in fine-tuning CaMKIIα activity and provide further support for the non-degradative roles of CaMKIIα ubiquitination. However, our proteomic data do not allow us to directly determine whether CaΜKIIα is mono- or polyubiquitinated at K291.

Previous studies have showed the regulation of kinase activities by intermolecular interactions with poly-ubiquitin chains (Kanayama et al., 2004; Xia et al., 2009; Yang et al., 2009). For instance, in the canonical NFκB signalling pathway, two kinases, TAK1 and IKK, undergo activation by interactions with K63-linked or K63/M1-linked ubiquitin chains^67,68,70^. However, the direct conjugation of ubiquitin to kinases, which serves non-degradative roles remains poorly understood (Filipčík et al., 2017). In this context, our study illustrates an emerging, additional facet of ubiquitin-mediated regulation through the direct conjugation of ubiquitin to a kinase.

## Materials & Methods

### Materials

LC/MS-grade water, acetonitrile (ACN), methanol, chloroform, CaCl2, KCl, KH2PO4, MgCl2, NaCl, NaHCO3, NaHPO4, 25% (v/v) NH4OH, sucrose and glucose were purchased from Merck, Darmstadt, Germany. Triethylammonium bicarbonate (TEAB), EGTA, PM400-Ficoll, NADP, L-glutamic dehydrogenase from bovine liver (GluDH), formic acid (FA), glycolic acid (GA), guanidine hydrochloride, tris(2-carboxylethyl)phosphine (TCEP), chloroacetamide (CAA), PR-619, ionomycin calcium salt were purchased from Sigma-Aldrich, Taufkirchen, Germany. HEPES was obtained from VWR Chemicals, Darmstadt, Germany. Trifluoracetic acid (TFA) was purchased from Roth, Karlsruhe, Germany. MS-grade trypsin and LysC were purchased from Promega, Madison, Wisconsin, USA. RapiGest was purchased from Waters, Milford, USA. Pierce 660 nm protein assay, Halt Protease and phosphatase inhibitor cocktail, and isobaric TMT6plex reagents (TMT6) were obtained from Thermo Fisher Scientific, Bleiswijk, Netherlands.

### Animals

All animals were handled in accordance with the regulations of the University of Göttingen and of the local authorities, the State of Lower Saxony (Landesamt für Verbraucherschutz, LAVES, Braunschweig, Germany). All animal experiments were performed in accordance with the European Communities Council Directive (2010/63/EU) and approved by the local authority, the Lower Saxony State Office for Consumer Protection and Food Safety (Niedersächsisches Landesamt für Verbraucherschutz und Lebensmittelsicherheit).

### Synaptosome preparation and glutamate release assay

Wistar rats aged between 5 and 7 weeks were sacrificed by cervical dislocation before decapitation. Whole brains were rapidly removed from the skull and cooled in ice-cold homogenisation buffer (320 mM sucrose, 5 mM HEPES, pH 7.4). Cerebral cortices and cerebella were subsequently dissected and homogenised at 900 rpm using 9 strokes with a Teflon/glass homogeniser. The resulting homogenate was used for the isolation of synaptosomes by the discontinuous Ficoll gradient centrifugation method, according to previously reported procedures (von Mollard et al., 1991). Specifically, the homogenate was centrifuged at 2988 *g* for 2 min and the resulting supernatant (S1) was subjected to an additional centrifugation step at 14462 *g* for 12 min using an SS-34 fixed-angle rotor (Thermo Fisher Scientific, Waltham, USA). The crude synaptosomal pellet (P2) was resuspended in ice-cold homogenisation buffer and layered onto a discontinuous Ficoll density gradient (6%/9%/13% w/v Ficoll in homogenisation buffer) followed by centrifugation at 86575 *g* for 40 min using an SW-41 swinging bucket rotor (Beckman Coulter, Krefeld, Germany). The synaptosome-enriched fraction at the interphase between 9% and 13% w/v Ficoll was obtained and washed with homogenisation buffer. Synaptosomes were concentrated in a final centrifugation step at 14462 *g* for 12 min using an SS-34 fixed-angle rotor (Thermo Fisher Scientific, Waltham, USA) and the resulting synaptosomal pellet was resuspended in ice-cold homogenisation buffer. Protein concentration was determined by the Pierce 660 nm protein assay following the manufacturer’s instructions.

The viability of synaptosomes was assessed by using the continuous fluorometric assay of glutamate release as previously described (Nicholls and Sihra, 1986). Briefly, 2–2.5 mg of synaptosomal protein was centrifuged at 6900 *g* for 3 min in a bench-top centrifuge and resuspended in 2 ml of physiological buffer (10 mM glucose, 5 mM KCl, 140 mM NaCl, 5 mM NaHCO3, 1 mM MgCl2, 1.2 mM NaHPO4, 20 mM HEPES, pH7.4). Synaptosomes were incubated in physiological buffer at 37 °C for 5 min to re-establish ATP levels, followed by the addition of 1 mM NADP and either 1.3 mM CaCl2 or 0.5 mM EGTA. Subsequently, synaptosomal suspension was transferred to a quartz glass cuvette (Hellma, Müllheim, Germany) and the incubation at 37 °C was continued with stirring for 3 more minutes. 200 u of glutamate dehydrogenase was further added to the synaptosomal suspension and the NADPH-fluorescence at 440 nm was measured for 3 min in a Fluorolog-3 fluorimeter (Horiba Jobin Yvon, Bensheim, Germany). 50 mM KCl was then added to the synaptosomal suspension to induce glutamate release, and the NADPH fluorescence was measured for a further 1 min. Finally, the synaptosomal suspension was centrifuged at 6900 *g* for 15 s in a bench-top centrifuge and the synaptosome pellet was lysed in 0.2 ml lysis buffer (6 M guanidine hydrochloride, 50 mM HEPES, pH 8, 10 mM TCEP, 40 mM CAA, 5 mM EDTA, 50 µM PR- 619, 1× Halt protease and phosphatase inhibitor cocktail).

### Protein precipitation and digestion

Lysed synaptosomes were incubated at 50 °C for 30 min in the presence of 10 mM TCEP/40 mM CAA to reduce disulphide bonds and carbamidomethylate cysteine residues, respectively. Samples were briefly cooled on ice and further sonicated for 10 min using 30 s on/30 s off iterations at maximum intensity in a Bioruptor ultrasonication device (Diagenode, Seraing, Belgium). Proteins were precipitated by the methanol/chloroform precipitation method (Wessel and Flügge, 1984): briefly, 4 and 1 sample volumes of ice-cold methanol and chloroform, respectively, were added to lysed synaptosomes, followed by the addition of 3× sample volume of water for phase separation. The samples were vortexed and subsequently centrifuged at 9000 *g* for 1 min at 4 °C. The upper phase was carefully discarded, and the aggregated proteins were washed with 1 ml of ice-cold methanol. Samples were vortexed and the proteins were precipitated by centrifugation at 16,000 *g* for 10 min at 4 °C. The resulting protein pellet was resuspended in a digestion buffer (0.1% RapiGest, 100 mM TEAB, pH 8) and sonicated for 10 min with 30 s on/30 s off iteration as described above. Proteins were pre-digested for 2 h at 37 °C with LysC at a protease-to-protein ratio of 1:300 (w/w). Finally, trypsin was added at a 1:90 (w/w) trypsin-to-protein ratio and the incubation was continued at 25 °C for 16 h. Next day, RapiGest was precipitated by acidifying the solution with 1% TFA and 1 h incubation at 37 °C. Peptide solutions were cleared in a bench-top centrifuge at maximum speed for 10 min and dried in a centrifugal Savant SpeedVac (Thermo Fisher Scientific, Waltham, USA). Finally, peptides were subjected to desalting using 50-mg Sep-Pak C18 Vac cartridges (Waters, Milford, Massachusetts) according to manufacturer’s instructions. An aliquot of the desalted peptides was taken for the proteome analysis and the desalted peptides were dried in a centrifugal Savant SpeedVac (Thermo Fisher Scientific, Waltham, USA).

### K-ε-GG peptide enrichment and TMT labelling

K-ε-GG peptide enrichment was performed using a K-ε-GG-specific antibody (PTM-Scan ubiquitin remnant motif K-ε-GG kit, Cell Signaling Technology, Kit#5562) chemically crosslinked to agarose beads as previously described (Udeshi et al., 2013). Briefly, 2–2.5 mg of dried peptides were dissolved in 1 ml of ice-cold IAP buffer (50 mM MOPS, pH 7.2, 50 mM NaCl, 10 mM Na3PO4) and sonicated in the Bioruptor for 5 min using a 30 s on/30 s off cycle. Peptide solutions were cleared by centrifugation in a bench-top centrifuge at maximum speed for 3 min at 4 °C. Cleared peptide solutions were incubated with 31.25 µg of K-ε-GG-specific antibody for 2 h at 4 °C with gentle end-over-end rotation. Ab-bead conjugates were then centrifuged at 2000 *g* for 1 min and the unbound fraction was carefully removed. Ab-bead conjugates were washed 3 times with ice-cold IAP buffer followed by two washes with 100 mM HEPES, pH 8. Thereafter, beads were centrifuged at 2000 *g* for 1 min and resuspended in 200 µl HEPES, pH 8. Chemical labelling of K-ε-GG peptides still bound to the antibody was performed by adding 400 µg of TMT6 isobaric reagents to each sample and incubating for 10 min at 25 °C with shaking at 1000 rpm (Udeshi et al., 2020). TMT labelling was then quenched with 0.025% (v/v) hydroxylamine for 5 min at 25 °C. After quenching, Ab-beads conjugates were combined and washed twice with 1× PBS buffer. Finally, K-ε-GG peptides were eluted by addition of 100 µl of 0.15% (v/v) TFA and incubation for 10 min at 25 °C followed by two repetitions of the addition and incubation steps. The resulting (K-ε-GG) peptides were cleaned using C18 spin columns (Harvard Apparatus, Holliston, USA) and dried in a centrifugal Savant SpeedVac (Thermo Fisher Scientific, Waltham, USA).

### K-ε-GG peptide enrichment for PRM analysis

For all targeted MS analysis (PRM), protein precipitation and digestion were carried out as described above (see Protein precipitation and digestion method section) with the following exceptions. 2 mg of synaptosomal proteins were precipitated using the methanol/chloroform precipitation method and subsequently digested with LysC and trypsin using a protease-to-protein ratio (w/w) of 1:200 and 1:50, respectively. K-ε-GG peptide enrichment was performed using the K-ε-GG-specific antibody (PTM-Scan HS Ubiquitin/SUMO remnant motif K-ε-GG kit, Cell Signaling Technology, Kit#59322) covalently linked to magnetic beads. For the validation of several sites 200 fmol of standard synthetic SpikeTideL K-ε-GG peptides (JPT Peptide Technologies, Berlin, Germany) were added to the endogenous peptide mixture before K-ε-GG peptide enrichment. For the absolute quantification of CaMKIIα peptides, the peptide mixture was split into two equal aliquots of 1 mg each and 350 fmol of standard AQUA QuantPro peptides (Thermo Fisher Scientific, Waltham, USA) were added to each aliquot before K-ε-GG peptide enrichment. Briefly, dried peptides were dissolved in 1 ml of ice-cold IAP Bind buffer and sonicated in a water bath for 2 min. Peptide solutions were cleared by centrifugation in a bench-top centrifuge at maximum speed for 5min at 4 °C. Peptide solutions were then incubated with 7 µl of K-ε-GG-specific antibody for 3 h at 4 °c with gentle end-over-end rotation. Ab-bead conjugates were then centrifuged at 2000 *g* for 5 s and the unbound fraction was carefully removed on a magnetic rack. Ab-bead conjugates were washed three times with ice-cold Wash Buffer followed by two washes with LC-MS grade water. Finally, K-ε-GG peptides were eluted by addition of 200 µl of 0.15% (v/v) TFA, 10% ACN and incubation for 15 min at 25 °C. The resulting K-ε-GG peptide solutions were dried in a centrifugal Savant SpeedVac prior to LC-MS/MS analysis. For the phosphatase treatment, dried K-ε-GG peptides were resuspended in 20 µl of 1x CutSmart buffer (New England Biolabs, B7204S) and incubated with 2.5 u of QuickCIP phosphatase (New England Biolabs, M0525L) at 37 °C for 1 h. After that, QuickCIP was heat inactivated at 80 °C for 2 min and the peptide solutions were desalted using C18 StageTips (Empore C18 SPE Disks, Sigma Aldrich, Darmstadt, Germany). Briefly, StageTips were conditioned by 100 µl of 100% MeCN followed by 100 µl of 80% ACN/0.1% TFA and by 3x 100 µl 2%ACN/0.1%TFA. Peptides were then loaded on StageTips, washed 3x with 100 ul of 2%ACN/0.1% TFA and eluted 2x with 50 µl of 50% ACN/0.1%TFA. Eluted peptides were dried using vacuum centrifugation prior to MS analysis.

### Phosphopeptide enrichment for PRM analysis

Phospho-peptide enrichment was performed with Zr-IMAC HP beads (MagReSyn, ReSyn Biosciences, Edenvale, South Africa). For this, beads were used in 4:1 beads-to-peptide ratio. Zr- IMAC HP beads are magnetic, therefore each supernatant removal step was executed by using a magnetic rack. Beads were equilibrated and washed with loading buffer (80% ACN, 5% TFA, 0.1 M glycolic acid). Dried, tryptic peptides were re-dissolved in loading buffer (3 min sonication) and added to the Zr-IMAC HP beads. Samples were then incubated at 25 °C for 30 min at 850 rpm. Subsequently, non-phospho-peptides were removed and either discarded or transferred to a fresh Eppendorf tube and dried. Beads were washed in three steps, first adding 500 µl of loading buffer with subsequent incubation of 1 min at 25 °C with 850 rpm, supernatant was removed and discarded and this step was repeated with wash buffer 1 (80% ACN, 1% TFA) and wash buffer 2 (10% ACN, 0.2% TFA). Phospho-peptides were eluted by adding 150 µl of 1% v/v ammonium hydroxide solution to the beads and 10 min incubation at 25 °C, 850 rpm. Phospho-peptides were then transferred to a fresh Eppendorf tube containing 50 µl 10% v/v TFA. A second elution step was performed in the same manner and the supernatant was added to the Eppendorf tube as well. Phospho-peptides were then dried in the SpeedVac (Thermo Fisher Scientific, Waltham, USA). For MS/MS-analysis, phospho-peptides were re-dissolved in 2% v/v ACN/0.05% v/v TFA.

### Basic reversed phase chromatography

For proteome analysis, TMT6-labeled peptides were separated by basic reversed-phase (bRP) chromatography with an Agilent 1100 series HPLC system (Agilent, Santa Clara, USA) equipped with a v C18-X-Bridge column (3.5 μm particles, 1.0 mm inner diameter, 150 mm length; Waters, Milford, USA). The HPLC was set to operate at a flow rate of 60 μl/min under basic conditions, with buffer A (10 mM NH4OH in water, pH ∼10) and buffer B (10 mM NH4OH and 80% (v/v) ACN in water, pH ∼10). The column was initially equilibrated with a mixture of 95% buffer A and 5% buffer B. A linear gradient ranging from 10% to 36% buffer B was then applied for 34 minutes, followed by a linear increase to 55% over 8 minutes and a wash step with 95% buffer B for 5 minutes. The resulting peptides were collected into 12 final fractions by concatenating one-minute fractions. Finally, the resulting bRP fractions were dried using a SpeedVac.

### LC-MS/MS analysis

Dried TMT-labelled peptides were resuspended in 5% (v/v) ACN, 0.1% (v/v) TFA in water and injected onto a C18 PepMap100-trapping column (0.3 × 5 mm, 5 µM, Thermo Fisher Scientific, Waltham, USA) coupled to a C18 analytical column packed in-house (75 µM × 300 mm, Reprosil- Pur 120C18- AQ, 1.9 µm, Dr Maisch, GmbH, Ammerbuch, Germany). The HPLC system was operated at a flow rate of 0.300 µl/min on an UltiMate-3000 RSLC nanosystem (Thermo Fisher Scientific, Waltham, USA). Both columns were equilibrated with a mixture of 95% buffer A (0.1% (v/v) FA in water) and 5% buffer B (80% (v/v) ACN, 0.1% (v/v) FA in water). TMT-labelled K-ε-GG peptides were eluted by using a linear gradient ranging from 14% to 38% buffer B for 90 min followed by a linear increase to 48% buffer B for 10 min and a wash step with 90% buffer B for 5 min. The eluted peptides were further injected into a QExactive HF-X (Thermo Fisher Scientific, Bremen, Germany), operated in data-dependent acquisition mode alternating between MS and MS2 acquisitions. TMT-labelled K-ε-GG peptides were analysed by using the following settings: MS1 scans in the range of 300-1400 *m/z* were acquired at a resolution of 120,000 at *m/z* 200, with an automatic gain control (AGC) of 10^6^ and a maximum injection time of 100 ms. The 20 most abundant precursor ions with charge state of +2 to +6 were selected using a 0.8 *m/z* isolation window and fragmented with a normalised collision energy (NCE) of 33%. MS2 fragment spectra were acquired with a resolution of 30,000, an AGC target of 10^5^ and a maximum injection time of 120 ms. Dynamic exclusion was applied for 20 s and a lock mass ion (*m/z* 445.1200) was used for internal calibration.

For the analysis of peptides not labelled with K-ε-GG TMT, similar settings were used, with the following exceptions: a linear gradient ranging from 10% to 36% buffer B for 62 min followed by a linear increase to 45% buffer B for 8 min was used. The eluted peptides were further injected into an Orbitrap Exploris 480 (Thermo Fisher Scientific, Bremen, Germany), where MS1 scans were acquired in the range of 300–1700 *m/z* and with a maximum injection time of 40 ms. The 30 most abundant precursor ions were selected using a 0.7 *m/z* isolation window and fragmented with a normalised collision energy (NCE) of 36%. MS2 fragment spectra were acquired with an AGC target of 5×10^4^ and a maximum injection time of 60 ms.

For the PRM analysis of K-ε-GG peptides, similar settings were used, with the following exceptions: a linear gradient ranging from 12% to 36% buffer B for 43 min followed by a linear increase to 45% buffer B for 3 min was used. The eluted peptides were further injected into an Orbitrap Exploris 480 (Thermo Fisher Scientific, Bremen, Germany), which was operated in targeted mass acquisition mode switching between MS1 and targeted MS2 scans. Within a 3-second cycle time one MS1 scan in the range of 350-1300 *m/z* was acquired with a maximum injection time of 50 ms followed by MS2 spectra derived from the targeted peptides. Specifically, heavy and light peptides matching the *m/z* values defined in the precursor isolation list (**Suppl. Data 2**) were isolated with a 1 *m/z* isolation window and fragmented with a normalised collision energy (NCE) of 28%. MS2 fragment spectra were acquired with a resolution of 60,000, an AGC target of 5×10^5^ and a maximum injection time of 120 ms.

### Peptide identification and data analysis

Raw files were analysed by using the MaxQuant (MQ) software (version 2.0.3.0) (Cox et al., 2011; Tyanova et al., 2016). Precursor ions and MS2 spectra were searched against the proteome of *Rattus norvegicus* containing canonical protein sequences (Uniprot (The UniProt Consortium, 2019), March 2021, 29,942 entries). Most of the MQ search settings were kept at default, with the following exceptions; apart from the default variable and fixed modifications (methionine oxidation, acetylation of protein N-termini and cysteine carbamidomethylation) K-ε-GG of lysine and TMT-6plex of lysine were set as variable modifications (only for the K-ε-GG peptides). Specific digestion with trypsin was selected allowing up to three missed cleavage sites per peptide. The maximum peptide mass was set to 5000. For K-ε-GG peptide quantification, reporter ions in MS2 level were selected with TMT6plex of the peptide N-termini as isobaric labels, whereas for non-K-ε-GG peptide quantification, TMT6plex values of the peptide N-termini and internal lysines were determined as isobaric labels. FDR at the PSM and protein levels were kept at the default setting of 1%.

All the downstream data analysis was performed in R statistical programming language using customised scripts. Impurity-corrected reporter ion intensities for each ubiquitination site were retrieved from the MaxQuant K-ε-GG site table. Ubiquitination sites assigned to potential contaminants, reversed sequences or with a localization probability lower then 0.75 as estimated by MQ were not considered in the downstream analysis. Ubiquitination sites with more than 3 zero values per TMT-6plex experiment were excluded from the quantification analysis. The remaining zero values (if any) were imputed using the minimal reporter ion intensity per channel. Corrected reporter ions were log2-transformed and normalised by using the Tukey median polishing procedure with a maximum iteration number of 3. Finally, statistical testing was conducted using the limma package (Ritchie et al., 2015).

Gene enrichment analysis was performed using the synapse-specific database, SynGO (SynGO release 20210225 ) (Koopmans et al., 2019) and ShinyGO (release version 0.77). Gene names of proteins associated with ubiquitination sites were used as foreground against a custom “synaptic proteome” background to extract significantly enriched (FDR corrected *p* value < 0.001) GO- biological process terms. An experimentally validated ubiquitination site data set downloaded from PhosphoSitePlus (Hornbeck et al., 2015) (February 2023) containing 105,710 unique ubiquitination sites was used as a literature reference and compared with our data set. Sequence windows of the ubiquitination sites in our data set were aligned against sequence windows in the PhosphoSitePlus data set with Blast software (Altschul et al., 1990) (version 2.13.0+), to account for possible sequence differences across different species.

### PRM data analysis

PRM raw data files were imported into Skyline (MacLean et al., 2010) (version 23.1.0.455) for manual inspection and refinement of integrated peak areas. Only peptides with at least 7 transitions were further considered for the quantification analysis. For quantification, low intensity transitions were not considered, and the peak areas of selected transitions were summed, as determined by Skyline (MacLean et al., 2010). Summed peak areas of endogenous peptides were normalised to summed peak areas of heavy peptides, as determined by Skyline (MacLean et al., 2010). Normalized peak area ratios were exported from Skyline and further subjected to statistical analysis in R.

### Cell culture and generation of CaMKIIα WT and CaMKIIα K291R HeLa Kyoto cell lines

HeLa Kyoto cells were cultured in high glucose Dulbecco’s modified Eagle’s medium (DMEM, Gibco, Thermo Fisher Scientific, Waltham, USA) supplemented with 10% foetal bovine serum (FBS), 2 mM glutamine, 1 mM sodium pyruvate and 100 units/ml penicillin, 0.1 mg/ml streptomycin at 37 °C and 5% CO2. Two different cell lines were generated: one cell line expressing wild-type CaMKIIα and another expressing the mutant variant CaMKIIα K291R. For this purpose, HeLa Kyoto cell lines were transfected with the Lipofectamine 3000 kit (Invitrogen, Thermo Fisher Scientific, Waltham, USA) and cells stably expressing the constructs 2 days after transfection were selected with 500 µg/ml geneticin (Gibco, Thermo Fisher Scientific, Waltham, USA).

### HeLa cell processing prior to PRM-MS analysis

HeLa cells stably exressing CaMKIIα were stimulated with 2.5 µM of the calcium ionophore, ionomycin in the presence of 1.8 mM CaCl2 in a cell incubator at 37 °C, and 5% CO2 for 7 min. Then, the medium was discarded and the cells were washed one time with 1x PBS buffer, followed by addition of lysis buffer (0.5% v/v NP40, 50 mM HEPES, pH 7.5, 100 mM NaCl, 1 mM EDTA; 50 µM PR-619, 40 mM CAA, 1× Halt protease and phosphatase inhibitor cocktail). Cells were scraped off and the cell lysates were transferred to tubes. Nuclei were pelleted by centrifuging at 10,000 *x g* for 30 s at 4°C in a bench-top centrifuge. The supernatants were transferred to new tubes and any remaining DNA was further digested by the Pierce universal nuclease (250 U) (Thermo Fisher Scientific, Waltham, USA) in the presence of 4 mM MgCl2 for 30 min at 37 °C. Subsequently, 20 mM TCEP/40 mM CAA were added to the solutions followed by incubation for 30 min at 37 °C to reduce disulphide bonds and carbamidomethylate cysteine residues. Afterwards, equal amounts of proteins as determined by BCA assay were cleaned up by the single-pot, solid-phase-enhanced sample-preparation (SP3) method as previously described (Cite Hughes et al., 2018) with a few modifications. Specifically, 10 mg of the bead stock (Sera-Mag Speedbeads, Cytiva;Marlborough, United States) were added per 1 mg of protein solution. To induce binding of the proteins to the beads 100% ACN was added to the solution to achieve a final ACN concentration of 50% v/v. The binding mixture was incubated at 24 °C for 5 min at 1,000 rpm. Afterwards, the tubes were placed in a magnetic rack and the unbound fraction was discarded. Afterwards, beads were washed three times with 80% v/v ethanol. A final washing step was performed with 100% v/v ACN. Beads were resuspended in digestion buffer (100 mM TEAB) and sonicated for 2 min in a water bath. Finally, proteins were digested for 16 h at 25 °C with trypsin at a trypsin-to-protein ratio of 1:30 (w/w). Next day, the tubes were placed to the magnetic rack and peptides solutions were collected to fresh tubes. 700 fmol of standard synthetic AQUA peptides per mg of protein amount (Technologies, Thermo Fisher Scientific, Waltham, USA) were added to the endogenous peptide mixture and peptides were dried in a centrifugal Savant SpeedVac (Thermo Fisher Scientific, Waltham, USA). Dried peptides were used either for K-GG peptide enrichment or phospho-peptide enrichment as previously described in method sections K-ε-GG peptide enrichment for PRM analysis and phosphopeptide enrichment for PRM analysis, respectively, prior to PRM-MS analysis (method section LC-MS/MS analysis).

### Preparation of rat dissociated hippocampal cultures

Newborn rats were used for the preparation of dissociated primary hippocampal cultures, as previously described (Kaech and Banker, 2006). In brief, hippocampi of newborn rat pups (wild-type, Wistar) were dissected in Hank’s Buffered Salt Solution (HBSS, 5 mM KCl, 140 mM NaCl, 4 mM NaHCO3, 0.3 mM Na2HPO4, 6 mM glucose and 0.4 mM KH2PO4). Subsequently, the tissues were incubated for one hour in enzyme solution (Dulbecco’s Modified Eagle Medium, DMEM, #D5671, Sigma-Aldrich, Germany), containing 0.5 mg/mL cysteine, 50 mM EDTA, 100 mM CaCl2 and 2.5 U/mL papain, saturated with carbogen for 10 min). The dissected hippocampi were then incubated for 15 min in a deactivating solution (DMEM with 0.2 mg/mL bovine serum albumin, BSA, 5% fetal calf serum and 0.2 mg/mL trypsin inhibitor). Then the cells were triturated and were seeded at a density of about 80,000 cells per 18 mm round coverslip. Before seeding, the coverslips underwent a treatment with nitric acid, then they were sterilized and coated overnight with 1 mg/mL poly-L-lysine. The cells were allowed to attach to the coverslips for a period between one to four hours at 37 °C in plating medium (DMEM with 10% horse serum, 2 mM glutamine and 3.3 mM glucose). The plating medium was then replaced with a Neurobasal-A medium (Life Technologies, Carlsbad, CA, USA) containing 1% GlutaMax (Gibco, Thermo Fisher Scientific, USA), 2% B27 (Gibco, Thermo Fisher Scientific, USA) supplement and 0.2% penicillin/streptomycin mixture (Biozym Scientific, Germany). Before use, the cultures were maintained in a cell incubator at 37 °C, and 5% CO2 for 9–11 days. Percentages represent volume/volume.

### Transfection of hippocampal neurons

Transfections were performed with a standard Lipofectamine 2000 kit (#11668019, ThermoFisher Scientific, Germany). In brief, neurons were pre-incubated for 25 min in 400 μl per well, pre-heated DMEM (#D5671, Sigma-Aldrich, Germany) complemented with 10 mM MgCl2 at pH 7.5 (fresh- DMEM). Per 18 mm coverslip, one 1 μg of DNA, prepared in a total volume of 25 μl Opti-MEM (#11058-021, Life Technologies Limited, United Kingdom) was used. The DNA-Opti-MEM solution was incubated for 5 min and added to 25 μl Opti-MEM with 1 μl lipofectamine solution. Then the solution was incubated for 15 min and subsequently added to the neurons. After incubating the neurons at 37 °C and 5% CO2 for 20 minutes, they were washed 2 times with fresh-DMEM, returned to their original culture medium and incubated at 37 °C and 5% CO2 until the conduction of the respective experiments.

### Live labeling and Fixation

Cells were incubated live with 1:200 monoclonal mouse anti-Synaptotagmin1 antibody, conjugated to ATTO647N (#105 311AT1, Synaptic Systems, Göttingen, Germany) for 60 min at 37 °C and 5% CO2. After 3 washes with Tyrode’s solution (124 mM NaCl, 30 mM glucose, 25 mM HEPES, 5 mM KCl, 1 mM MgCl2, 2 mM CaCl2, pH 7.4), the cells were fixed in 4% PFA in PBS (10 mM Na2HPO4, 137 mM NaCl, 2. 7 mM KCl, 2 mM KH2PO4, pH 7.4) for 20 min at room temperature. The fixation reaction was quenched with 100 mM NH4Cl in PBS for 20 min. For subsequent immunostainings, neurons were permeabilized and blocked with PBS containing 0.1% Triton X (#9005-64-5, Merck, Germany), 5%, bovine serum albumin (BSA) (#A1391-0250; Applichem, Germany) for 30 min.

### Imaging and image analysis

The neurons were imaged with an inverted Nikon Ti microscope (Nikon Corporation, Chiyoda, Tokyo, Japan) with a Plan Apochromat 60x objective (1.4 NA, immersion oil). For the image analysis, a custom-made Matlab (Matlab version 2022b, the Mathworks Inc., Natick, MA, USA) macro was used. Briefly, synapses were identified based on the Syt1 signal. The fluorescence signal of Syt1 in the synaptic boutons was correlated with the CaMKIIα expression signal within the area of each synapse, using Pearson correlation. Subsequently, the fluorescence intensity of Syt1 was quantified in the boutons in which the Syt1 and the CaMKIIα signals correlated well. A paired t-test between the wild type and the mutant was performed to determine significant differences (p=0.03).

## Acknowledgements

We thank Reinhard Jahn for his support in this study. We further thank the Facility of Light Microscopy, and especially Peter Lenart, Antonio Politi and Jasmin Jakobi for their assistance in HeLa cell culture, FACS (fluorescence-activated cell sorting) and live imaging. We further acknowledge Ralf Pflanz, Monika Raabe, Uwe Plessmann, Olexandr Dybkov and Sabine König for help with MS, as well as Sascha Krause, Thomas Gundlach and Ulrike Teichmann for their assistance in animal experiments.

## Competing interests

The authors state that they have no conflicts of interest with the contents of the article.

## Funding

H.U, S.O.R. were funded by the Deutsche Forschungsgemeinschaft and the collaborative research center SFB1286 (project numbers A08 and A03). S.L. was funded by Deutsche Forschungsgemeinschaft and the collaborative research center SFB1565 (project number P17).

## Author contributions

H.U., I.S.., and S.A. conceived the study; S.A designed and conducted synaptosome experiments under the supervision of I.S. and H.U; S.A. performed MS data analysis with the guidance of I.S. and H.U.; S.L.., and S.A. conceptualized the stimulation experiments of CaMKIIα in HeLa (Kyoto) cells; S.A. generated the mutant CaMKIIα, generated stably transfected HeLa cell lines and performed the stimulation experiments of Hela cells with the guidance of S.L; S.K. performed the phosphopeptide enrichment; S.K. and S.A. performed the PRM and statistical analysis in HeLa cells CaMKIIα system; S.O.R and S.G. conceptualized the synaptic activity assay of neurons expressing CaMKIIα and S.G. performed the experiments; S.A., H.U. wrote the manuscript with input from the other co-authors.

## Notes

### Competing Interest Statement

The authors have declared no competing interest.

