## Supplementary Material for "Calcium-triggered (de)ubiquitination events in synapses"

**Supplementary figures:**

### 1.1. Supplementary figures

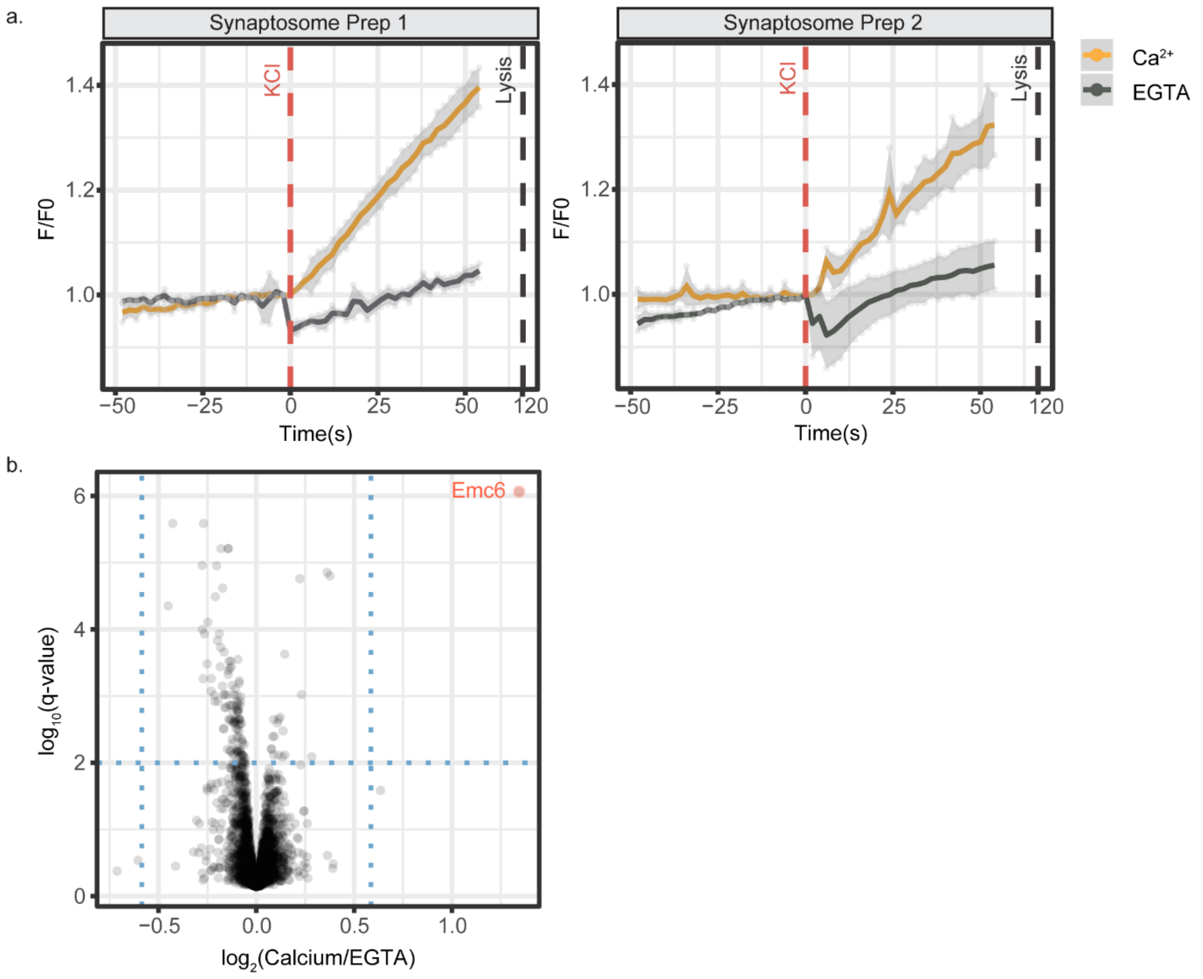

***Figure S1: Glutamate release assay and proteome changes in depolarized synaptosomes under different stimuli.*** *a) KCl depolarization of synaptosomes in the presence of $Ca^{2+}$ or EGTA as a control to account for $Ca^{2+}$-independent exocytosis. Glutamate release was monitored in response to chemical depolarization in real time (x-axis) and correlated with normalized fluorescence intensity (F/F0) (y-axis). Yellow line corresponds to the average response of synaptosomes depolarized in the presence of $Ca^{2+}$ from three independent stimulation experiments. Grey line corresponds to the average response of synaptosomes depolarized in the presence of EGTA from three independent stimulation experiments. Shaded grey areas represent the standard deviation at each time point .b) Volcano plot showing $log_2$(intensity fold change) of proteins quantified under $Ca^{2+}$ vs. EGTA conditions against $-log_{10}$(q-value). Proteins that show at least an abundance change of 1.5-fold at 1% FDR are coloured in red.*

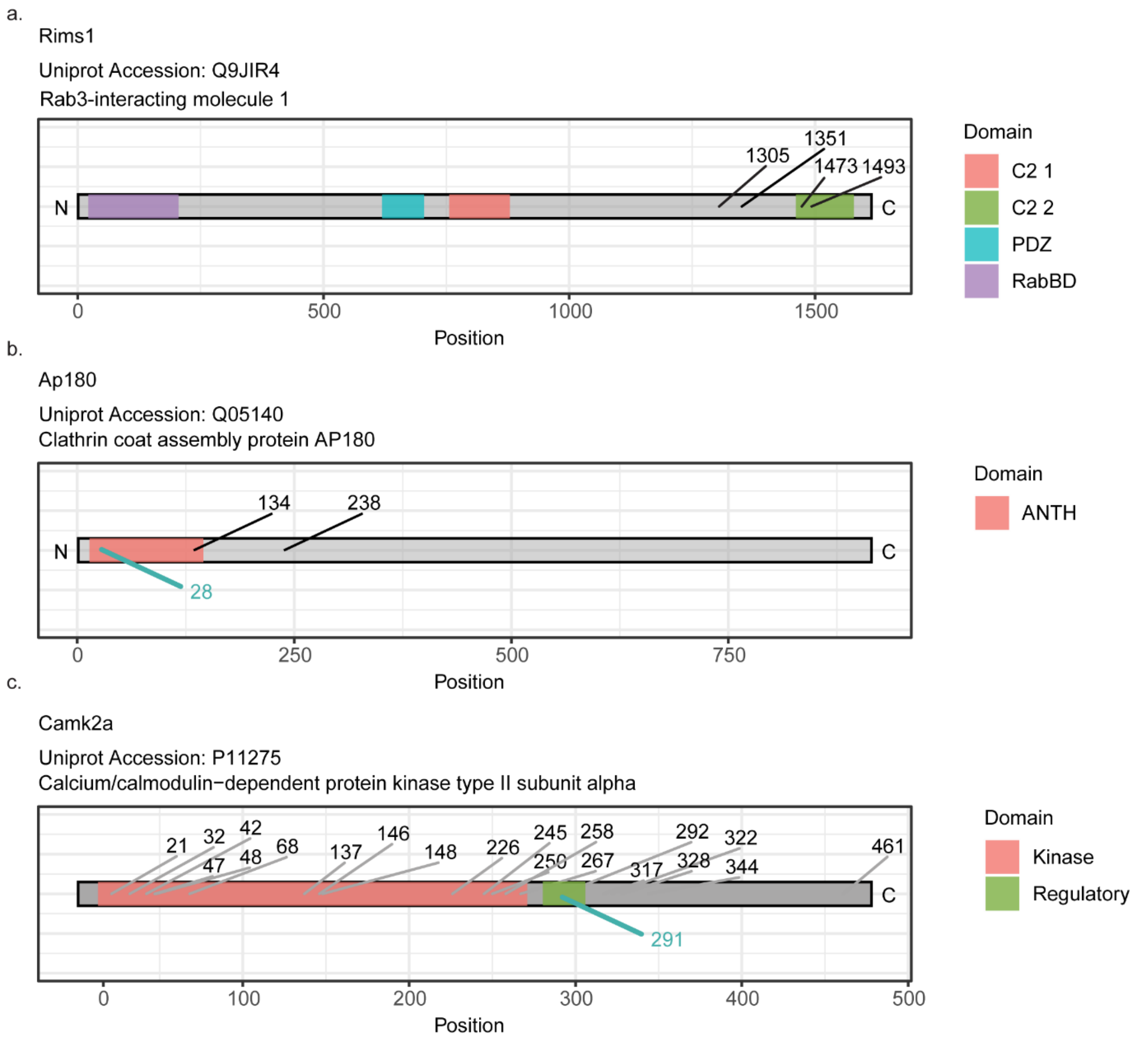

***Figure S2: Ubiquitination sites mapped at selected proteins.*** *The protein sequence is shown as a grey bar and the corresponding domains are color-coded. The exact position of the ubiquitination sites with PEP <0.01 as determined by MaxQuant*[1,2] *are shown. The ubiquitination sites not affected by* $Ca^{2+}$ *influx are shown in black, whereas those sites that show abundance changes in response to* $Ca^{2+}$ *influx are highlighted in cyan blue. a, b) The domains of Rims1 and Ap180 are presented as annotated in UniProt*[3]*, whereas c) the domains of CaMKIIα are manually annotated according to Chao et al.*[4]*.*

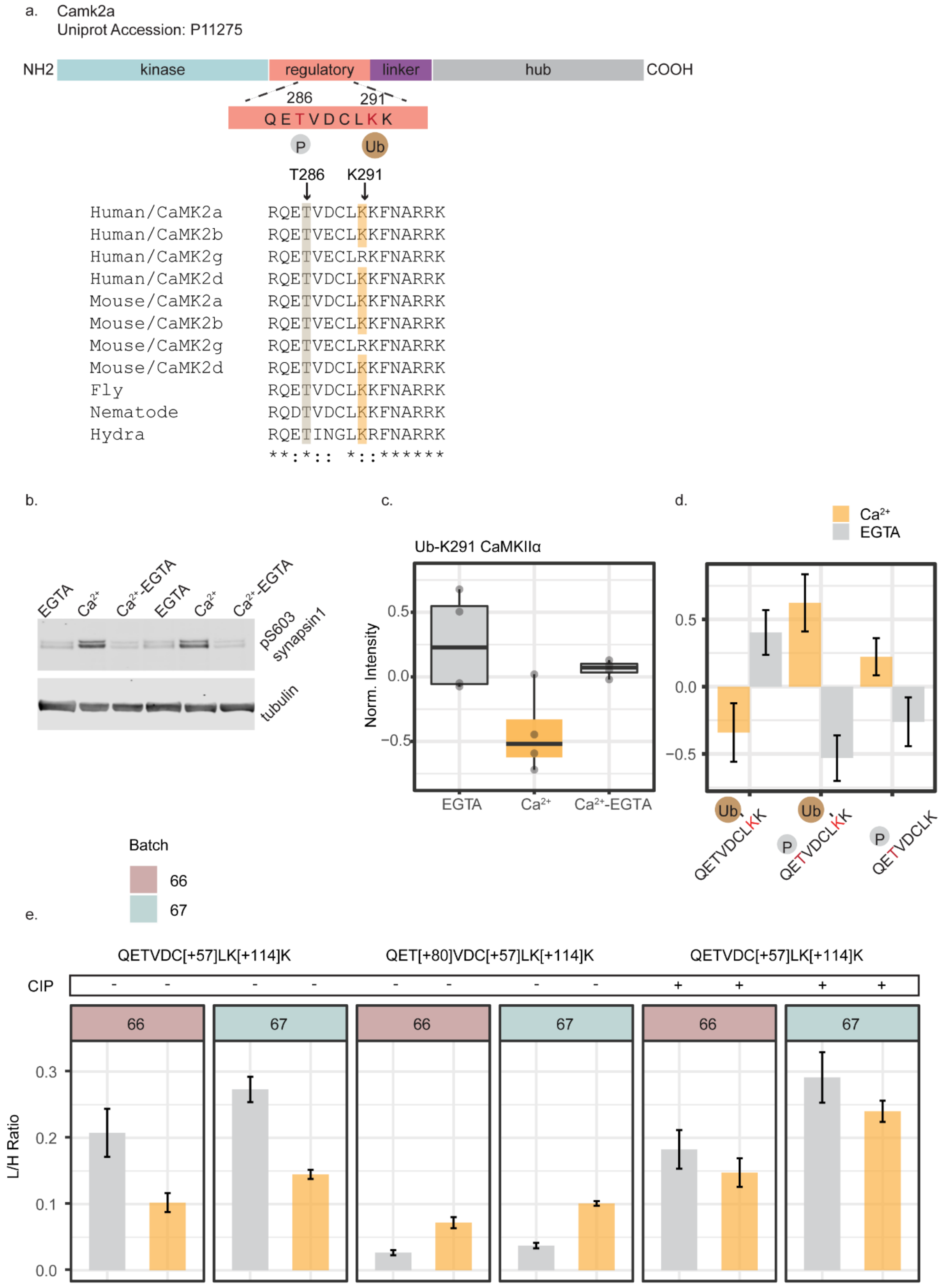
a.
Camk2a
Uniprot Accession: P11275
NH2
kinase
regulatory
linker
hub
COOH
286
291
QETVDCLKK
P
Ub
T286
K291
Human/CaMK2a RQETVDCLKKFNARRK
Human/CaMK2b RQETVECLKKFNARRK
Human/CaMK2g RQETVECLRKFNARRK
Human/CaMK2d RQETVDCLKKFNARRK
Mouse/CaMK2a RQETVDCLKKFNARRK
Mouse/CaMK2b RQETVECLKKFNARRK
Mouse/CaMK2g RQETVECLRKFNARRK
Mouse/CaMK2d RQETVDCLKKFNARRK
Fly RQETVDCLKKFNARRK
Nematode RQDTVDCLKKFNARRK
Hydra RQETINGLKRFNARRK
**:*:: *::******
b.
EGTA
Ca2+
Ca2+-EGTA
EGTA
Ca2+
Ca2+-EGTA
pS603 synapsin1
tubulin
c.
Ub-K291 CaMKIIα
Norm. Intensity
0.5
0.0
−0.5
EGTA
Ca2+
Ca2+-EGTA
d.
Ca2+
EGTA
0.5
0.0
−0.5
Ub
QETVDCLKK
Ub
P
QETVDCLKK
P
QETVDCLK
e.
Batch
66
67
QETVDC[+57]LK[+114]K
QET[+80]VDC[+57]LK[+114]K
QETVDC[+57]LK[+114]K
CIP
- - - - - - - - + + + +
66
67
66
67
66
67
L/H Ratio
0.3
0.2
0.1
0.0

***Figure S3: Post-translational modifications mapped at the regulatory domain of CaMKIIα and their abundance changes in depolarized synaptosomes under different stimuli.*** a) *Protein sequence alignment of the regulatory domain of CaMKII homologues was performed using the multiple sequence alignment tool T-Coffee[5]. The regulatory T286 and K291 (numbering based on human CaMKIIα with UniprotID Q9UQM7) are highlighted with grey and yellow colours, respectively. b) SDS/PAGE and immunoblotting against phosphosynapsin and tubulin (as a control for gel loading) were performed. An increase in the phosphorylation state of synapsin was observed in the presence of $Ca^{2+}$, indicative of functional synaptosomes. Upon $Ca^{2+}$ chelation the phosphorylation level of synapsin decreased and returned to basal levels. (c) A bottom-up proteomic workflow was used for the TMT-based quantification of formerly ubiquitinated (K-ε-GG) peptides. Boxplot shows K-ε-GG modified peptide corresponding to ubiquitinated CaMKIIα at K291. The x-axis corresponds to the different treatments of the synaptosomes and the y-axis corresponds to the normalized TMT reporter ion intensities. d) Peptide species corresponding to part of the regulatory region of CaMKIIα were detected either ubiquitinated on K291 or doubly modified by ubiquitination and phosphorylation under our experimental conditions. The figure shows log2-normalised reporter-ion intensities of differently modified peptide species derived from two independent TMT6 experiments. The phosphorylated peptide species corresponding to autophosphorylated CaMKIIα at T286 was included in the plot based on our previous quantification of the phosphorylation sites in resting and excited synaptosomes[6]. e) Barplot showing the mean light-to-heavy peak area ratios of modified CaMKIIα peptides before and after QuickCIP treatment from two different batches of synaptosomal preparation. Error bars correspond to the standard estimation estimated by three independent stimulation replicates.*

### 1.2. Supplementary tables

***Table S1: Enriched GO biological function terms of all identified ubiquitinated proteins according to SynGO database:*** *A list of proteins with well-localized ubiquitination sites identified in Ca-vs-EGTA experiments was used as a foreground in pathway enrichment analysis using the SynGO database. GO biological functions terms that were significantly enriched at an enrichment FDR <0.01 are shown.*

| GO term ID | GO term name - hierarchical structure | FDR |
|---|---|---|
| SYNGO:synprocess | process in the synapse | 1.54E-14 |
| SYNGO:presynprocess | ├─ process in the presynapse | 3.78E-10 |
| GO:0099504 | │ ├─ synaptic vesicle cycle | 4.97E-08 |
| GO:0016079 | │ │ ├─ synaptic vesicle exocytosis | 0.0047 |
| GO:0048488 | │ │ └─ synaptic vesicle endocytosis | 0.002181 |
| SYNGO:postsynprocess | ├─ process in the postsynapse | 5.02E-05 |
| GO:0099072 | │ ├─ regulation of postsynaptic membrane neurotransmitter receptor levels | 0.001806 |
| GO:0099536 | ├─ synaptic signaling | 0.0047 |
| GO:0099537 | │ ├─ trans-synaptic signaling | 0.0047 |
| GO:0050808 | ├─ synapse organization | 0.002993 |

***Table S2: Enriched GO biological function terms of regulated ubiquitinated proteins according to SynGO database:*** *A list of proteins with regulated ubiquitination sites as determined in Ca-vs-EGTA experiments was used as a foreground in pathway enrichment analysis using the SynGO database. GO biological functions terms that were significantly enriched at an enrichment FDR <0.01 are shown.*

| GO term ID | GO term name - hierarchical structure | FDR |
|---|---|---|
| SYNGO:synprocess | process in the synapse | 8.9972E-06 |
| SYNGO:presynprocess | ├─ process in the presynapse | 1.45029E-07 |
| GO:0099504 | │ ├─ synaptic vesicle cycle | 3.50589E-07 |
| GO:0016079 | │ │ ├─ synaptic vesicle exocytosis | 8.9972E-06 |
| GO:0048488 | │ │ └─ synaptic vesicle endocytosis | 0.000366676 |
| GO:0099525 | │ ├─ presynaptic dense core vesicle exocytosis | 1.00844E-05 |
| SYNGO:postsynprocess | ├─ process in the postsynapse | 0.005995807 |

***Table S3: Enriched GO biological function terms of all identified ubiquitinated proteins according to ShinyGO database*** *A list of proteins with well-localized ubiquitination sites identified in Ca-vs-EGTA experiments was used as a foreground in pathway enrichment analysis using the ShinyGO database. GO biological functions terms that were significantly enriched at an enrichment FDR <0.005 are shown.*

| Pathway | Enrichment FDR | Fold Enrichment |
|---|---|---|
| Synaptic vesicle cycle | 5.19E-10 | 1.958419 |
| Vesicle-mediated transport in synapse | 1.06E-09 | 1.888838 |
| Regulation of neurotransmitter levels | 3.71E-08 | 1.854714 |
| Neurotransmitter transport | 1.88E-07 | 1.828719 |
| Regulated exocytosis | 2.59E-07 | 1.807157 |
| Neurotransmitter secretion | 4.97E-07 | 1.907698 |
| Signal release from synapse | 4.97E-07 | 1.907698 |
| Potassium ion transport | 1.43E-06 | 1.8335 |
| Regulation of exocytosis | 4.07E-06 | 1.79824 |
| Synaptic vesicle exocytosis | 3.72E-05 | 1.932437 |
| Potassium ion transmembrane transport | 9.29E-05 | 1.785838 |
| Regulation of neurotransmitter transport | 0.000118115 | 1.933845 |
| Postsynapse organization | 0.000140545 | 1.703963 |
| Regulation of neuronal synaptic plasticity | 0.000293933 | 2.2478 |
| Regulation of synaptic plasticity | 0.000342866 | 1.6405 |
| Negative regulation of transmembrane transport | 0.001226744 | 1.918123 |
| Regulation of neurotransmitter secretion | 0.001226744 | 1.883248 |
| Regulation of proteasomal protein catabolic process | 0.001475548 | 1.686625 |
| Cytokinesis | 0.001733313 | 1.79824 |
| Cell-cell junction assembly | 0.001733313 | 1.961716 |
| Adenylate cyclase-modulating G protein-coupled receptor signaling pathway | 0.001733313 | 1.835445 |
| Negative regulation of ion transmembrane transport | 0.001733313 | 1.986627 |
| Negative regulation of ion transport | 0.001733313 | 1.849618 |

| | | |
|---|---|---|
| Regulation of proteasomal ubiquitin-dependent protein catabolic process | 0.002787001 | 1.804336 |
| Glycolytic process | 0.002791192 | 1.991897 |
| Negative regulation of cation transmembrane transport | 0.002791192 | 1.991897 |
| ATP generation from ADP | 0.002791192 | 1.991897 |
| Synaptic vesicle recycling | 0.002791192 | 1.964906 |
| Negative regulation of neuron death | 0.002839177 | 1.663914 |
| Regulation of regulated secretory pathway | 0.002839177 | 1.72631 |
| Import across plasma membrane | 0.003132779 | 1.743748 |
| Negative regulation of neuron apoptotic process | 0.004875467 | 1.717722 |
| Cytoskeleton-dependent cytokinesis | 0.004875467 | 1.918123 |
| Regulation of proteolysis involved in cellular protein catabolic process | 0.004937502 | 1.585387 |
| Sodium ion transmembrane transport | 0.004937502 | 1.72631 |

***Table S4: Selected K-ε-GG peptides for targeted PRM analysis in depolarized synaptosomes:*** *26 K-ε-GG peptides representing 26 ubiquitination sites were selected for PRM analysis. The exact modified peptide sequence of the targets is presented; [+57] and [+114] correspond to the nominal masses of carbamidomethylation and K-ε-GG ubiquitin remnant, respectively.*

| **Protein** | **Position** | **Uniprot ID** | **Peptide sequence** |
|---|---|---|---|
| CaMK2a | K291 | P11275 | QETVDC[+57]LK[+114]K |
| AP180 | K28 | Q05140 | AVC[+57]K[+114]ATTHEVMGPK |
| Unc13c | K992 | Q62770 | ILAGDSSSVDEK[+114]AR |
| Caps1 | K953 | Q62717 | TDYNLC[+57]NGK[+114]FHK |
| Caps1 | K1249 | Q62717 | LQGVLDSTLNSK[+114]TYETIR |
| Syt7 | K375 | Q62747 | IYLSWK[+114]SGPGEVK |
| Stx1b | K93 | P61265 | LK[+114]AIEQSIEQEEGLNR |
| Stx1a | K94 | P32851 | LK[+114]SIEQSIEQEEGLNR |
| Stx4 | K102 | Q08850 | AQLK[+114]AIEPQK |
| Snap25 | K76 | P60881 | EAEK[+114]NLTDLGK |
| Snap25 | K103 | P60881 | K[+114]AWGNNQDGVVASQPAR |
| Vamp2 | K59 | P63045 | DQK[+114]LSELDDR |
| Vamp7 | K172 | Q9JHW5 | TENLVDSSVTFK[+114]TTSR |
| Ppap2b | K15 | P97544 | AIVPESK[+114]NGGSPALNNNPR |
| Pclo | K2766 | D3Z9C7 | LHSYVK[+114]AEEDPMEDPYELK |
| Calm2 | K31 | P0DP31 | DGDGTITTK[+114]ELGTVMR |
| Usp24 | K645 | F1LSM0 | SFIK[+114]QTYQK |
| Camk2b | K292 | F1LNI8 | QETVEC[+57]LK[+114]K |
| Usp5 | K423 | D3ZVQ0 | ALIGK[+114]GHPEFSTNR |
| Ubc | K6 | P62982 | MQIFVK[+114]TLTGK |
| Ubc | K11 | P62982 | TLTGK[+114]TITLEVEPSDTIENVK |
| Ubc | K27 | P62982 | TITLEVEPSDTIENVK[+114]AK |
| Ubc | K29 | P62982 | AK[+114]IQDKEGIPPDQQR |
| Ubc | K33 | P62982 | IQDK[+114]EGIPPDQQR |
| Ubc | K48 | P62982 | LIFAGK[+114]QLEDGR |
| Ubc | K63 | P62982 | TLSDYNIQK[+114]ESTLHLVLR |

***Table S5: Selected peptides for the absolute quantification analysis of CaMKIIα in synaptosomes and stimulated HeLa (Kyoto) cells:*** *Eight peptides derived from CaMKIIα were monitored in depolarized synaptosomes and stimulated HeLa (Kyoto) cells. The exact modified peptide sequence of the targets is presented; [+57], [+80], and [+114] correspond to the nominal masses of carbamidomethylation, phopshorylation and K-ε-GG ubiquitin remnant, respectively. * refers to peptides targeted exclusively in the HeLa cells but not in the depolarized synaptosomes.*

| Protein | Peptide sequence |
|---|---|
| CaMK2a | ITAAEALK |
| | ITQYLDAGGIPR |
| | QET[+80]VDC[+57]LK |
| | QET[+80]VDC[+57]LK[+114]K |
| | QETVDC[+57]LK |
| | QETVDC[+57]LK[+114]K |
| | RITAAEALK |
| | VLAGQEYAAK |
| CaMK2a Mut* | QET[+80]VDC[+57]LR* |
| | QETVDC[+57]LR* |

***Table S6: Ubiquitinated E3 ligases in depolarized synaptosomes***

| Uniprot ID | Gene name | Protein description | Sites |
|---|---|---|---|
| Q5PQN2 | Bfar | Bifunctional apoptosis regulator | 387 |
| Q8CIN9 | Rffl | E3 ubiquitin-protein ligase rififylin | 258 |
| Q66H79 | Trim32 | Tripartite motif protein 32 | 508;407;473;361 |
| D3ZL82 | Trim37 | Tripartite motif protein 37 (Predicted) | 411 |
| D3ZJB8 | Arih2 | RBR-type E3 ubiquitin transferase | 376 |
| D3ZXL1 | Arih1 | RBR-type E3 ubiquitin transferase | 142;437 |
| Q5XIQ4 | Mgrn1 | E3 ubiquitin-protein ligase MGRN1 | 165;102 |
| F1M8V2 | Ube4b | Ubiquitination factor E4B | 665 |
| A0A0G2JVW5 | Huwe1 | HECT-type E3 ubiquitin transferase | 2698;324 |
| Q62940 | Nedd4 | E3 ubiquitin-protein ligase NEDD4 | 556;450;483 |
| F1LRN8 | Nedd4l | HECT-type E3 ubiquitin transferase | 456;624;506;524;447 |

***Table S 7: Ubiquitinated DUBs in depolarized synaptosomes.***

| Uniprot ID | Gene name | Protein description | Sites |
|---|---|---|---|
| B2GUZ1 | Usp4 | Ubiquitin carboxyl-terminal hydrolase 4 | 835 |
| D3ZPG5 | Usp30 | Ubiquitin carboxyl-terminal hydrolase 30 | 338;310 |
| Q4VSI4 | Usp7 | Ubiquitin carboxyl-terminal hydrolase 7 | 870;421 |
| Q6J1Y9 | Usp19 | Ubiquitin carboxyl-terminal hydrolase 19 | 836;843 |
| Q5D006 | Usp11 | Ubiquitin carboxyl-terminal hydrolase 11 | 819;794 |
| D3ZC84 | Usp9x | Ubiquitinyl hydrolase 1 | 1627;1632;1811;112;1798;315;1435 |
| F1LPJ7 | Usp33 | Ubiquitin carboxyl-terminal hydrolase | 226 |
| D3ZLQ8 | Usp20 | Ubiquitin carboxyl-terminal hydrolase | 325;907 |

| | | | |
|---|---|---|---|
| D3ZVQ0 | Usp5 | Ubiquitin carboxyl-terminal hydrolase | 423;476;743;508;574;184;64;318;357;360;391;406;178;793;558;20 |
| D4ACD3 | Usp25 | Ubiquitin specific protease 25 (Predicted) | 251;631 |
| A0A0G2JTF5 | Usp34 | Ubiquitinyl hydrolase 1 | 1317 |
| A0A0G2JZU8 | Usp9y | Ubiquitin-specific peptidase 9, Y-linked | 1809 |
| D3ZBB7 | Usp32 | Ubiquitinyl hydrolase 1 | 810 |
| F1LSM0 | Usp24 | Ubiquitin-specific peptidase 24 | 418;1761;650;645 |
| Q5U2N2 | Usp14 | Ubiquitin carboxyl-terminal hydrolase | 448;342;238;300;214 |
| Q00981 | Uchl1 | Ubiquitin carboxyl-terminal hydrolase isozyme L1 | 71;115;123;4 |
| O35815 | Atxn3 | Ataxin-3 | 200 |
| Q8CF97 | Vcpip1 | Deubiquitinating protein VCPIP1 | 181;385;763;710;690;697;1012;375;361;878;198;407;330;457;140 |
| O88767 | Park7 | Parkinson disease protein 7 homolog | 182;93;130;41 |
| Q499N6 | Ubxn1 | UBX domain-containing protein 1 | 178;83;105 |
| B2RYG6 | Otub1 | Ubiquitin thioesterase OTUB1 | 188;71;73 |
| Q32Q05 | Yod1 | Ubiquitin thioesterase OTU1 | 252 |
| D3ZH40 | Otud7b | Ubiquitinyl hydrolase 1 | 263;93 |
| Q66HP6 | Spata2 | Spermatogenesis-associated protein 2 | 27 |
| Q8R424 | Stambp | STAM-binding protein | 107 |
| Q5BJY4 | Josd1 | Josephin-1 | 180 |
| D4ABZ4 | Otud7a | Ubiquitinyl hydrolase 1 | 89 |
| P46462 | Vcp | Transitional endoplasmic reticulum ATPase | 696;236;8;18;486;60;251;231;668;651;217;502;658 |
| B2RYM5 | Brcc3 | Lys-63-specific deubiquitinase BRCC36 | 180 |

### 1.3. Supplementary data

***Suppl. Data 1:*** *TMT reporter ion intensities used for the assessment of regulated ubiquitination site in EGTA-vs-Ca2+ treated synaptosomes.*

***Suppl. Data 2:*** *Method parameters of PRM analysis*

***Suppl. Data 3:*** *Light to heavy ratios of targeted peptides used for the assessment of regulated ubiquitination sites in EGTA -vs- Ca2+ treated synaptosomes by PRM.*

***Suppl. Data 4:*** *Light to heavy ratios of targeted CaMKIIa peptides used for the assessment of changes in K291 ubiquitination and T286 phosphorylation of CaMKIIα in EGTA -vs- Ca2+ treated synaptosomes by PRM.*

***Suppl. Data 5:*** *Light to heavy ratios of targeted peptides used for the assessment of changes in K291 ubiquitination and T286 phosphorylation of CaMKIIα in DMSO-vs-ionomycin treated HeLa (Kyoto) cells.*
